# Intratumoral delivery of engineered recombinant modified vaccinia virus Ankara expressing Flt3L and OX40L generates potent antitumor immunity through activating the cGAS/STING pathway and depleting tumor-infiltrating regulatory T cells

**DOI:** 10.1101/2021.10.31.466698

**Authors:** Ning Yang, Yi Wang, Shuaitong Liu, Joseph M. Luna, Gregory Mazo, Adrian Y. Tan, Tuo Zhang, Jiahu Wang, Wei Yan, John Choi, Anthony Rossi, Jenny Zhaoying Xiang, Charles M. Rice, Taha Merghoub, Jedd D. Wolchok, Liang Deng

## Abstract

Intratumoral (IT) delivery of immune-activating viruses can serve as an important strategy to turn “cold” tumors into “hot” tumors, resulting in overcoming resistance to immune checkpoint blockade (ICB). Modified vaccinia virus Ankara (MVA) is a highly attenuated, non-replicative vaccinia virus that has a long history of human use. Here we report that IT recombinant MVA (rMVA), lacking E5R encoding an inhibitor of the DNA sensor cyclic GMP-AMP synthase (cGAS), expressing a dendritic cell growth factor, Fms-like tyrosine kinase 3 ligand (Flt3L), and a T cell co-stimulator, OX40L, generates strong antitumor immunity, which is dependent on CD8^+^ T cells, the cGAS/STING-mediated cytosolic DNA-sensing pathway, and STAT1/STAT2-mediated type I IFN signaling. Remarkably, IT rMVA depletes OX40^hi^ regulatory T cells via OX40L/OX40 interaction and IFNAR signaling. Taken together, our study provides a proof-of-concept for improving MVA-based cancer immunotherapy, through modulation of both innate and adaptive immunity.

**One Sentence Summary:** Intratumoral delivery of recombinant MVA for cancer immunotherapy

## Introduction

Immune checkpoint blockade (ICB) therapy utilizing antibodies targeting T cell inhibitory mechanisms has revolutionized how solid tumors are treated (Ribas and Wolchok, 2018; Wei et al., 2018; Zou et al., 2016). However, the majority of patients without pre-existing antitumor T cell responses do not respond to ICB therapy and one-third of the initial responders develop acquired resistance to this line of therapy likely due to cancer immuno-editing (Ribas and Wolchok, 2018; Schreiber et al., 2011; Zaretsky et al., 2016). Therefore, innovative approaches to rendering tumors sensitive to ICB therapy are urgently needed.

Viral-based cancer immunotherapy is a versatile and effective approach to alter tumor immunosuppressive microenvironment (TME) through multiple mechanisms, including the induction of innate immunity, immunogenic cell death in the infected immune and tumor cells, and activation of tumor-infiltrating dendritic cells (DCs), and antitumor CD8 and CD4 T cells, as well as depletion of immunosuppressive cells (Bommareddy et al., 2018; Davola and Mossman, 2019; Lemos de Matos et al., 2020; Russell et al., 2012; Workenhe and Mossman, 2014). As a result, intratumoral (IT) delivery of immunogenic viruses turns “cold” tumors into “hot” tumors, which renders them sensitive to other immunotherapeutic modalities including ICB (Chesney et al., 2018; Dai et al., 2017; Ribas et al., 2018; Wang et al., 2021; Zamarin et al., 2014).

Poxviruses are large cytoplasmic DNA viruses. Modified vaccinia virus Ankara (MVA) is a highly attenuated vaccinia virus that has been used extensively as a vaccine vector (Gilbert, 2013; Liu et al., 2021; Volz and Sutter, 2017). MVA infection of dendritic cells induces type I IFN via the cytosolic DNA-sensing pathway mediated by the DNA sensor cyclic GMP-AMP synthase (cGAS) and downstream signaling molecules such as STimulator of INterferon Genes (STING) (Dai et al., 2014). However, MVA encodes multiple inhibitors of the nucleic acid-sensing pathways. IT heat-inactivated MVA (Heat-iMVA) generates stronger antitumor immunity than IT live MVA, which requires CD8^+^ T cells, Batf3-dependent CD103^+^/CD8*α* cross-presenting dendritic cells (DCs), and STING-mediated cytosolic DNA-sensing pathway (Dai et al., 2017).

We designed our recombinant MVA virus to engage both innate and adaptive immunity in the tumor microenvironment. First, given the importance of the cGAS/STING pathway in innate immune-sensing of MVA and MVA-induced antitumor immunity (Dai et al., 2017; Deng et al., 2014; Woo et al., 2014), we deleted the E5R gene, encoding a cGAS inhibitor, from the MVA genome to generate MVAΔE5R, which induces much higher levels of type I IFN compared with MVA (Yang et al., 2021). Second, we engineered the virus to express two membrane-anchored transgenes, FMS-like tyrosine kinase 3 ligand (Flt3L) and OX40L. Flt3L is a growth factor for CD103^+^ DCs and plasmacytoid DCs (Liu and Nussenzweig, 2010). OX40L is a co-stimulatory ligand for OX40, a member of the tumor necrosis factor (TNF) receptor superfamily expressed on activated CD4 and CD8 T cells as well as regulatory T cells (Croft, 2009). Here we used rMVA to designate MVAΔE5R-hFlt3L-mOX40L, which expresses human Flt3L and murine OX40L, and rhMVA to designate MVAΔE5R-hFlt3L-hOX40L, which expresses human Flt3L and human OX40L.

We observed that IT rMVA induces strong antitumor effects via the cGAS/STING-mediated DNA-sensing mechanism and the IFNAR/STAT1/STAT2 pathway. Depletion of CD8^+^ T cells renders tumors resistant to rMVA therapy. IT rMVA dramatically reduces OX40^hi^ regulatory T cells (Tregs) in the injected tumors via OX40L/OX40 interaction and IFNAR signaling. Taken together, our study strongly supports that rational engineering of MVA is an innovative strategy to deplete intratumoral OX40^hi^ Tregs to enhance antitumor immunity.

## Results

### The rationale for engineering a recombinant MVA virus (rMVA) with deletion of E5R and expression of human FMS-like tyrosine kinase 3 ligand (hFlt3L) and murine OX40L (mOX40L) for cancer immunotherapy

Our previous work demonstrated that Batf3-dependent CD103^+^ DCs are required for antitumor immunity induced by intratumoral (IT) delivery of heat-inactivated modified vaccinia virus Ankara (Dai et al., 2017). Flt3L is a growth factor that is important for DC development, especially for CD103^+^ DCs and plasmacytoid DCs (Liu and Nussenzweig, 2010). To investigate whether human Flt3L (hFlt3L) expression on tumor cells affects tumor growth and tumor-infiltrating myeloid cell populations, we constructed a murine melanoma B16-F10 stable cell line that expresses membrane-bound hFlt3L, and subsequently implanted either B16-F10-hFlt3L or the parental B16-F10 cells into WT C57BL/6J mice (**Figure S1A**). We observed that expressing hFlt3L on tumor cells delayed B16-F10 tumor growth and prolonged the survival of tumor-bearing mice (**Figure S1B and S1C**). The percentages of CD103^+^ DCs out of CD45^+^ cells, as well as the absolute numbers of CD103^+^ DCs per gram of B16-F10hFlt3L, were increased in B16-F10-hFlt3L tumors compared with B16-F10 control tumors, whereas CD11b^+^ DCs were at similar levels in both tumors (**Figure S1D**). These results indicate that hFlt3L expression on tumor cell surfaces facilitates the development and proliferation of CD103^+^ DCs in the tumor microenvironment.

OX40L is a costimulatory molecule that interacts with its receptor OX40 expressed on T cells (Croft et al., 2009). OX40L on activated dendritic cells plays an important role in the generation of antigen-specific T cell responses (Murata et al., 2000). We constructed a B16-F10-mOX40L cell line that constitutively expresses murine OX40L on its surface (**Figure S1A**). B16-F10-mOX40L tumors grew slower than the parental B16-F10 tumors after implantation (**Figure S1B and S1C**), with higher percentages of Granzyme B^+^ CD8^+^ T cells compared with the parental B16-F10 cells (**Figure S1E**). The median survival of mice implanted with B16-F10-hFlt3L or B16-F10-mOX40L were 28 and 34 days, 8 or 14 days longer, respectively, than those implanted with the control B16-F10 (**Figure S1C).**

We recently discovered that the vaccinia E5R gene encodes a potent inhibitor of cGAS (Yang et al., 2021). MVAΔE5R infection of murine bone marrow-derived dendritic cells (BMDCs) induces much higher levels of IFNB compared with MVA (**Figure S2A**). We compared the antitumor efficacy of MVA vs. MVAΔE5R in the B16-F10 melanoma unilateral implantation model in vivo (**Figure S2B**) and found that IT MVA prolonged the medium survival from 11 days in the PBS-treated group to 26 days, and IT MVAΔE5R resulted in 60% survival (**Figure S2C)**. We also designed the following recombinant MVA viruses to evaluate the utility of hFlt3L and mOX40L individually expressed by MVAΔE5R (**Figure S2D**). MVAΔE5R-hFlt3L and MVAΔE5R-mOX40L expressed respective transgenes in infected BHK21 cells (**Figure S2E**). IT delivery of the two viruses resulted in higher numbers of IFN-*γ*^+^ T cells in the spleens compared with MVA or MVAΔE5R, as determined by ELISPOT analysis (**Figure S2F and S2G**). These results suggest that expressing hFlt3L or mOX40L by recombinant MVA improves antitumor efficacy.

Based on these results, we designed an rMVA (MVAΔE5R-hFlt3L-mOX40L) by inserting two transgenes, a membrane-bound hFlt3L, and mOX40L into the E5R locus (**Figure 1A**). The hFlt3L and mOX40L are linked by a P2A self-cleaving sequence and their expression is driven by the vaccinia synthetic early/late promoter. We observed that both transgenes were expressed efficiently on the surface of infected B16-F10 murine melanoma cells and BMDCs at 24 h post-infection (**Figure 1B**). rMVA infection of BMDCs induced the expression of *Ifnb, Ifna4, Ccl4, Ccl5, Cxcl9, Cxcl10*, and *Il12p40* genes. Infection of BMDCs with rMVA induced cGAS-dependent *Ifnb* gene expression and IFN-*β* protein secretion at higher levels compared with MVA (**Figure 1C and 1D**). rMVA infection also induces DC maturation, as manifested by CD86 upregulation, determined by FACS, in a cGAS-dependent manner (**Figure 1E**).

**Figure 1.**
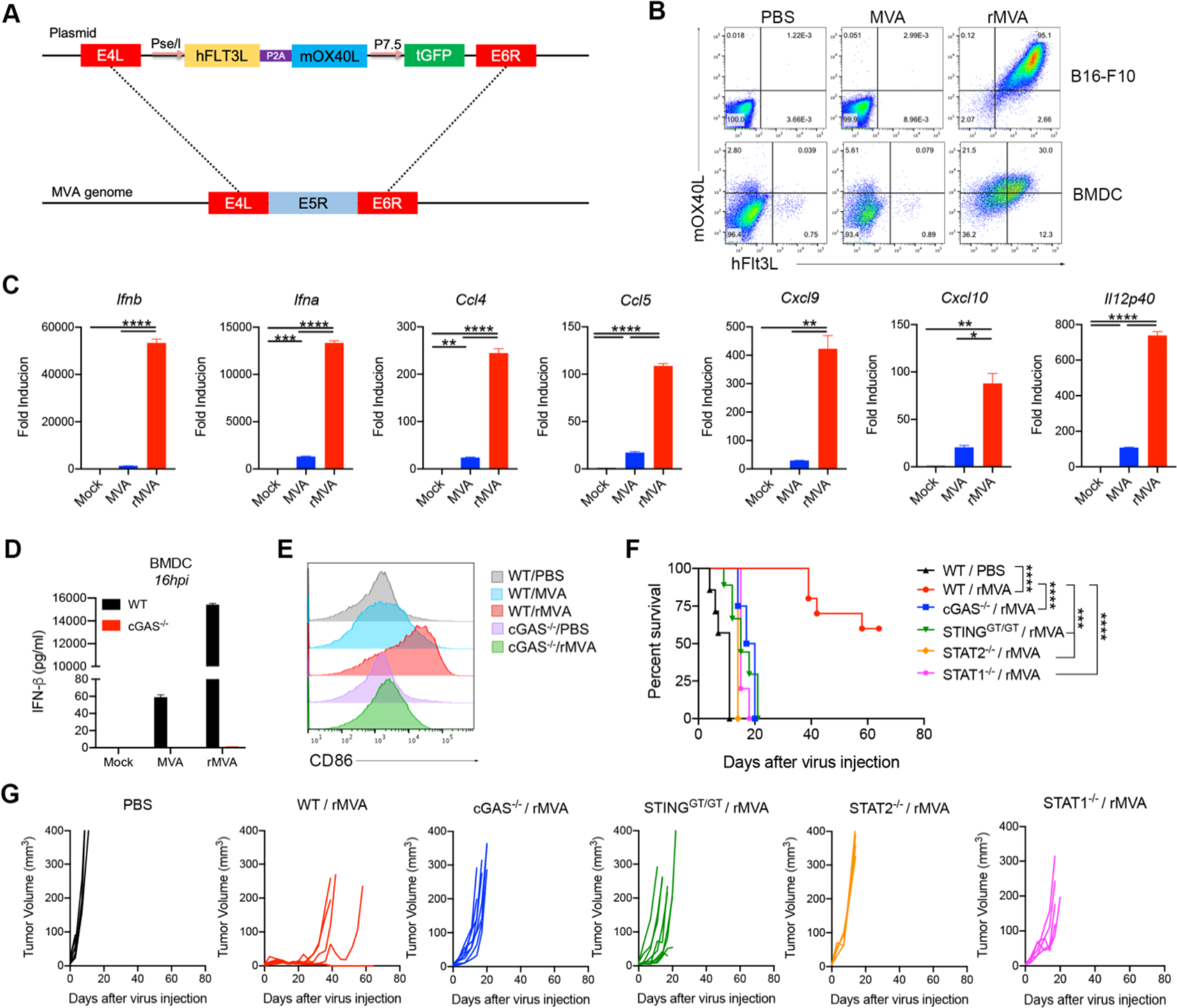
Intratumoral injection (IT) of rMVA elicits strong antitumor immune responses that is dependent on cGAS-STING and STAT1/STAT2. (A) Schematic diagram for the generation of rMVA through homologous recombination. (B) Representative flow cytometry plots of expression of hFlt3L or mOX40L by rMVA-infected B16-F10 cells and BMDCs. (C) Relative mRNA expression levels of *Ifnb*, *Ifna*, *Ccl4*, *Ccl5*, *Cxcl9, Cxcl10* and *Il12p40* in BMDCs infected with MVA or rMVA. Data are means ± SD (*n=3; *P < 0.05, **P < 0.01, ***P < 0.001, ****P < 0.0001, t test)*. (D) Concentrations of secreted IFN-*β* in the medium of WT or cGAS^-/-^ BMDCs infected with MVA or rMVA. Data are means ± SD. (E) Mean fluorescence intensity of CD86 expressed by WT or cGAS^-/-^ BMDCs infected with MVA or rMVA. (F) Kaplan-Meier survival curve of mice treated with rMVA or PBS in a unilateral B16-F10 implantation model (*n=5∼10; ***P < 0.001, ****P < 0.0001, Mantel-Cox test*). (G) Tumor growth curve of mice treated with rMVA or PBS in a unilateral B16-F10 implantation model.

### Intratumoral injection (IT) of rMVA elicits strong antitumor immune responses that are dependent on cGAS-STING-mediated DNA sensing and STAT1/STAT2-mediated IFNAR-signaling pathway

To test whether cGAS/STING and STAT1/STAT2 are important for IT rMVA-induced antitumor immunity, cGAS^-/-^, STING^Gt/Gt^ (lacking function STING) (Sauer et al., 2011), STAT1^-/-^, STAT2^-/-^ or age-matched WT C57BL/6J mice were implanted with B16-F10 melanoma intradermally. When the tumors were established, they were injected with rMVA twice weekly. Whereas IT rMVA resulted in tumor eradication or delayed tumor growth in WT mice, neither STAT1^-/-^ or STAT2^-/-^ mice responded to this therapy, with median survival of 15 and 12 days, respectively, compared with 11 days in the mock-treated group (**Figure 1F and 1G**). IT rMVA treatment of cGAS^-/-^ or STING^Gt/Gt^ mice extended median survival from 11 days in PBS control group to 18.5 days (*p* = 0.0002). However, all of the cGAS^-/-^ or STING^Gt/Gt^ mice died from tumor progression (**Figure 1F and 1G**). These results demonstrated that activation of the cGAS-mediated cytosolic DNA-sensing pathway, as well as the IFNAR/STAT1/STAT2 signaling, by IT rMVA, is critical for the generation of antitumor immunity.

### IT rMVA results in myeloid cells influx into the injected tumors and induces IFN-β and other inflammatory cytokine production in a cGAS/STING-dependent manner

To elucidate mechanisms of action of rMVA, we first investigated myeloid cell dynamics and determined which cell types are infected after IT viral therapy. To do that, we used the murine B16-F10 tumor implantation model and injected the tumors with MVAΔE5R expressing mCherry. At one or two-days post-injection, we harvested the tumors and analyzed tumor-infiltrating immune cells. IT MVAΔE5R-mCherry injection led to an influx of neutrophils one day post-injection, which subsided after the second day, when monocytes started to increase in the tumor microenvironment (**Figure 2A**). Among the myeloid cell populations, macrophages were most heavily infected, as determined by mCherry expression in the infected cells, followed by monocytes, neutrophils, CD103^+^, and CD11b^+^ DCs (**Figure 2B-2G**). T, B, or NK cells, however, were largely not infected by IT injected MVAΔE5R (**Figure 2G**).

**Figure 2.**
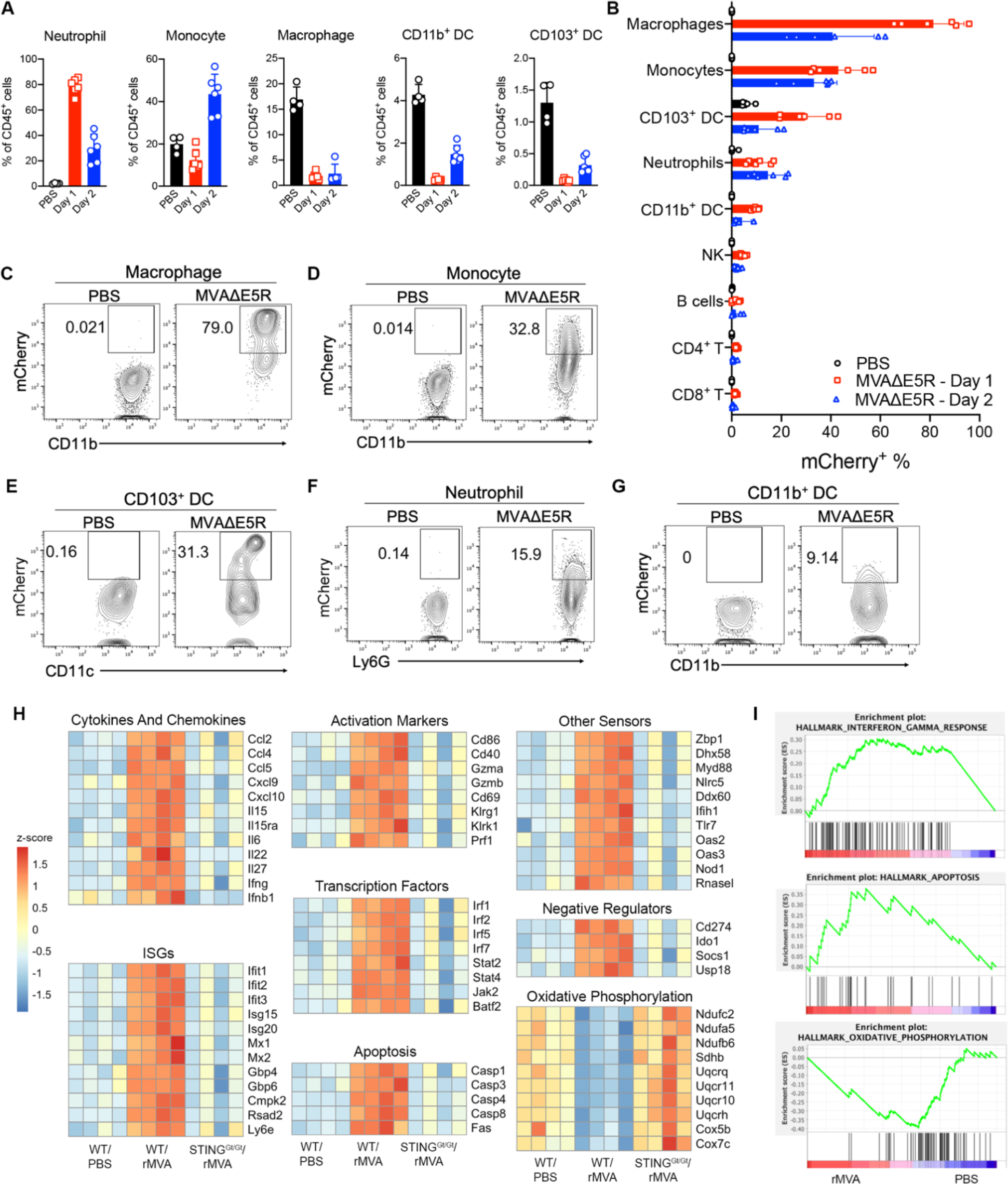
Influx of myeloid cells into MVAE5R-treated tumors and induction of IFN-β and other inflammatory cytokine production in a cGAS/STING-dependent manner. (A) Percentages of neutrophils, monocytes, macrophages, CD103^+^ DCs and CD11b^+^ DCs in the MVAΔE5R-treated tumors. Mice were intradermally implanted with B16-F10 cells. 7 days post implantation, tumors were injected with MVA*Δ*E5R-mCherry or PBS as control and harvested 1- or 2-days post injection for myeloid cell analysis. Data are means ± SD (*n=4∼6)*. (B) Percentages of mCherry^+^ immune cells. Data are means ± SD (*n=4∼6).* (C-G) Representative flow cytometry plots of mCherry^+^ immune cells. (H) Heatmap of gene expression from bulk tumor RNA-seq analysis. (I) Gene set enrichment analysis.

To investigate the innate immune responses of tumors induced by IT rMVA and the role of the cytosolic DNA-sensing pathway in this process, we isolated tumors from WT or STING^Gt/Gt^ mice one day post treatment with rMVA or PBS (mock control) and subjected them to bulk RNA-seq analyses. We observed striking upregulation of genes involved in immune activation, apoptosis, and downregulation of genes involved in oxidative phosphorylation (**Figure 2H and 2I**). We separated the immune activation genes into several subcategories, including cytokines and chemokines, interferon-stimulated genes (ISGs), activation markers, transcription factors, and other sensors. We found that rMVA treatment upregulated the expression of *Ifnb1*, *Ifng*, *Il-15*, *Il15ra*, *Ccl2*, *Ccl4*, *Ccl5*, *Cxcl9*, and *Cxcl10* in a STING-dependent manner (**Figure 2H and 2I**). We also observed up-regulation of DC activation markers, including *CD86* and *CD40*, and T cell activation markers, including *Gzma*, *Gzmb*, *Pfr1*, and *CD69,* as dependent on STING (**Figure 2H and 2I**). *Caspase 1*, *Caspase 3*, *Caspase 4*, *Caspase 8,* and *Fas* gene expression was also induced by IT rMVA in WT mice but not in STING-deficient mice (**Figure 2H and 2I**). The expression of genes involved in oxidative phosphorylation was downregulated after IT rMVA injection in WT mice but not in STING-deficient mice (**Figure 2H and 2I**). Taken together, our results show that IT rMVA leads to infection and recruitment of myeloid cell populations and activation of innate immune responses in those cells via the cGAS/STING pathway.

### IT rMVA generates stronger systemic and local anti-tumor immune responses compared with MVAΔE5R in a B16-F10 murine melanoma implantation model

To determine the immunological mechanism of rMVA-induced antitumor immune responses, we used a murine B16-F10 bilateral tumor implantation model. B16-F10 cells were intradermally implanted into both flanks of C57BL/6J mice. After tumors were established, we injected MVAΔE5R, rMVA, or PBS as a control, to the right-side tumors twice, three days apart. Spleens and both tumors were harvested 2 days after the second injection (**Figure 3A**). IT rMVA generated the highest numbers of tumor-specific IFN-γ^+^ T cells in the spleens compared with those treated with MVAΔE5R or with PBS as determined by ELISpot assay (**Figure 3B and 3C**). In the injected tumors, IT rMVA resulted in stronger T cell activation with higher percentages and absolute numbers of granzyme B^+^ CD8^+^ and CD4^+^ cells compared with MVAΔE5R, or PBS control groups (**Figure 3D-3F**). In the non-injected tumors, IT rMVA also induced more granzyme B^+^ CD8^+^ and CD4^+^ T cells (**Figure 3G-3I**), demonstarting that IT rMVA enhances T cell activation both locally and systemically. IT rMVA also induced IFN-γ^+^ TNF-α^+^ CD8^+^ and CD4^+^ T cells in the injected tumors, indicating enhanced T cell effector function (**Figure 3J-3M**). Taken together, these results demonstrate that IT rMVA results in the activation of both CD8^+^ and CD4^+^ T cells in the injected and non-injected tumors, as well as in the generation of systemic antitumor immunity.

**Figure 3.**
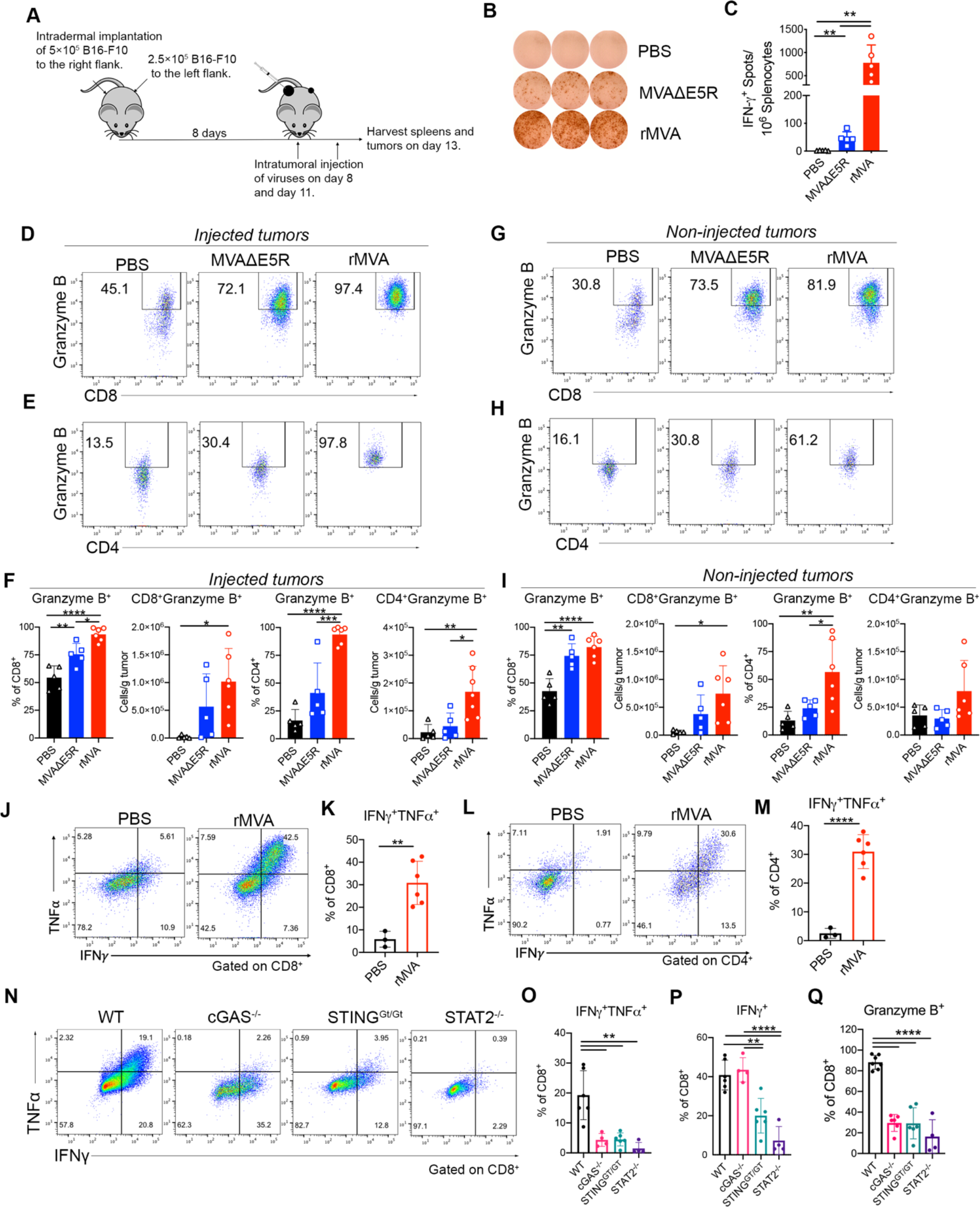
IT rMVA generates strong systemic and local anti-tumor immune responses which is dependent on cGAS/STING/STAT2 pathways. (A) Schematic diagram of IT rMVA or MVA*Δ*E5R for ELISpot assay and TIL analysis in a murine B16-F10 melanoma implantation model. (B) Representative images of IFN-*γ*^+^ spots from ELISpot assay. (C) Statistical analysis of IFN-*γ*^+^ splenocytes from MVA*Δ*E5R, rMVA or PBS-treated mice. Data are means ± SD (*n=5 or 6; **P < 0.01, t test)*. (D-E) Representative flow cytometry plots of Granzyme B^+^ CD8^+^ (D) and Granzyme B^+^ CD4^+^ cells (E) in the injected tumors. (F) Percentages and absolute number of Granzyme B^+^ CD8^+^ and Granzyme B^+^ CD4^+^ cells in the injected tumors. Data are means ± SD (*n=5 or 6; *P < 0.05, **P < 0.01, ***P<0.001, ****P < 0.0001, t test)*. (G-H) Representative flow cytometry plots of Granzyme B^+^ CD8^+^ (G) and Granzyme B^+^ CD4^+^ cells (H) in the non-injected tumors. (I) Percentages and absolute number of Granzyme B^+^ CD8^+^ (J) and Granzyme B^+^ CD4^+^ (K) cells in the non-injected tumors. Data are means ± SD (*n=5 or 6; *P < 0.05, **P < 0.01, **** P < 0.0001, t test)*. (J-M) Representative flow cytometry plots and statistical analysis of IFN*γ*^+^TNF*α*^+^ CD8^+^ (J, K) and IFN*γ*^+^TNF*α*^+^ CD4^+^ cells (L, M) in the injected tumors. Data are means ± SD (*n=3 or 6; **P < 0.01, ****P < 0.0001, t test)*. (N) Representative flow cytometry plots of IFN*γ*^+^TNF*α*^+^ CD8^+^ T cells in the injected tumors harvested from WT, cGAS^-/-^, STING^Gt/Gt^, and STAT2^-/-^ mice. (O) Percentages of IFN*γ*^+^TNF*α*^+^ CD8^+^ T cells in the injected tumors from WT, cGAS^-/-^, STING^Gt/Gt^ and STAT2^-/-^ mice. Data are means ± SD (*n=4∼6; **P < 0.01, t test)*. (P) Percentages of IFN*γ*^+^ CD8^+^ T cells in the injected tumors from WT, cGAS^-/-^, STING^Gt/Gt^ and STAT2^-/-^ mice. Data are means ± SD (*n=4∼6; **P < 0.01, t test)*. (Q) Percentages of Granzyme B^+^ CD8^+^ T cells in the injected tumors from WT, cGAS^-/-^, STING^Gt/Gt^ and STAT2^-/-^ mice. Data are means ± SD (*n=4∼6; ****P < 0.0001, t test)*.

### rMVA-induced antitumor immune responses are dependent on the cGAS/STING-mediated cytosolic DNA-sensing and STAT2-mediated type I IFN signaling pathways

We observed that the B16-F10-bearing cGAS, STING, or STAT2-deficient mice responded poorly to rMVA treatment (**Figure 1K and 1L**), which led us to hypothesize that the cGAS/STING-mediated cytosolic DNA-sensing and STAT2-dependent IFN signaling pathways are important for the generation of antitumor CD8^+^ T cell responses. To test that hypothesis, cGAS, STING, or STAT2-deficient and age-matched C57BL/6J control mice were intradermally implanted with B16-F10 cells into their right flanks. IT rMVA generated polyfunctional IFN-γ^+^ TNF-α^+^CD8^+^ and granzyme B^+^ CD8^+^ T cells in the injected tumors only in WT mice, whereas only IFN-γ^+^ CD8^+^ T cells were induced by IT rMVA in cGAS^-/-^ and STING^Gt/Gt^ mice (**Figure 3N-3Q**). IT rMVA failed to induce either IFN-γ^+^, Granzyme B^+^ CD8^+^ cells, or IFN-γ^+^ TNF-α^+^ T cells in STAT2^-/-^ mice (**Figure 3N-3Q**). These results indicate that both the cGAS/STING and STAT2-mediated signaling pathways are crucial for rMVA-induced T cell activation.

### IT rMVA depletes OX40^hi^ Tregs in the injected tumors

In addition to enhanced CD8^+^ and CD4^+^ T cell activation, we also observed that IT rMVA treatment resulted in a significant reduction of Tregs in the injected tumors (**Figure 4A and 4B**). The mean percentages of Tregs (Foxp3^+^CD4^+^) out of CD4^+^ T cells were 23%, 45%, and 51% in rMVA, MVAΔE5R, or PBS-treated tumors, respectively (**Figure 4A and 4B**). The absolute numbers of Tregs in rMVA injected tumors were significantly reduced compared with the PBS-treated group (**Figure 4A and 4B**). In the non-injected tumors, however, we did not observe a reduction of the percentages of Tregs out of CD4^+^ T cells after rMVA treatment (**Figure 4C**). The percentages of cleaved caspase-3^+^ cells out of tumor-infiltrating Tregs from rMVA-treated tumors were much higher compared with those from PBS-treated tumors (**Figure 4D and 4E**). These results support that rMVA treatment triggers apoptosis in tumor-infiltrating Tregs.

**Figure 4.**
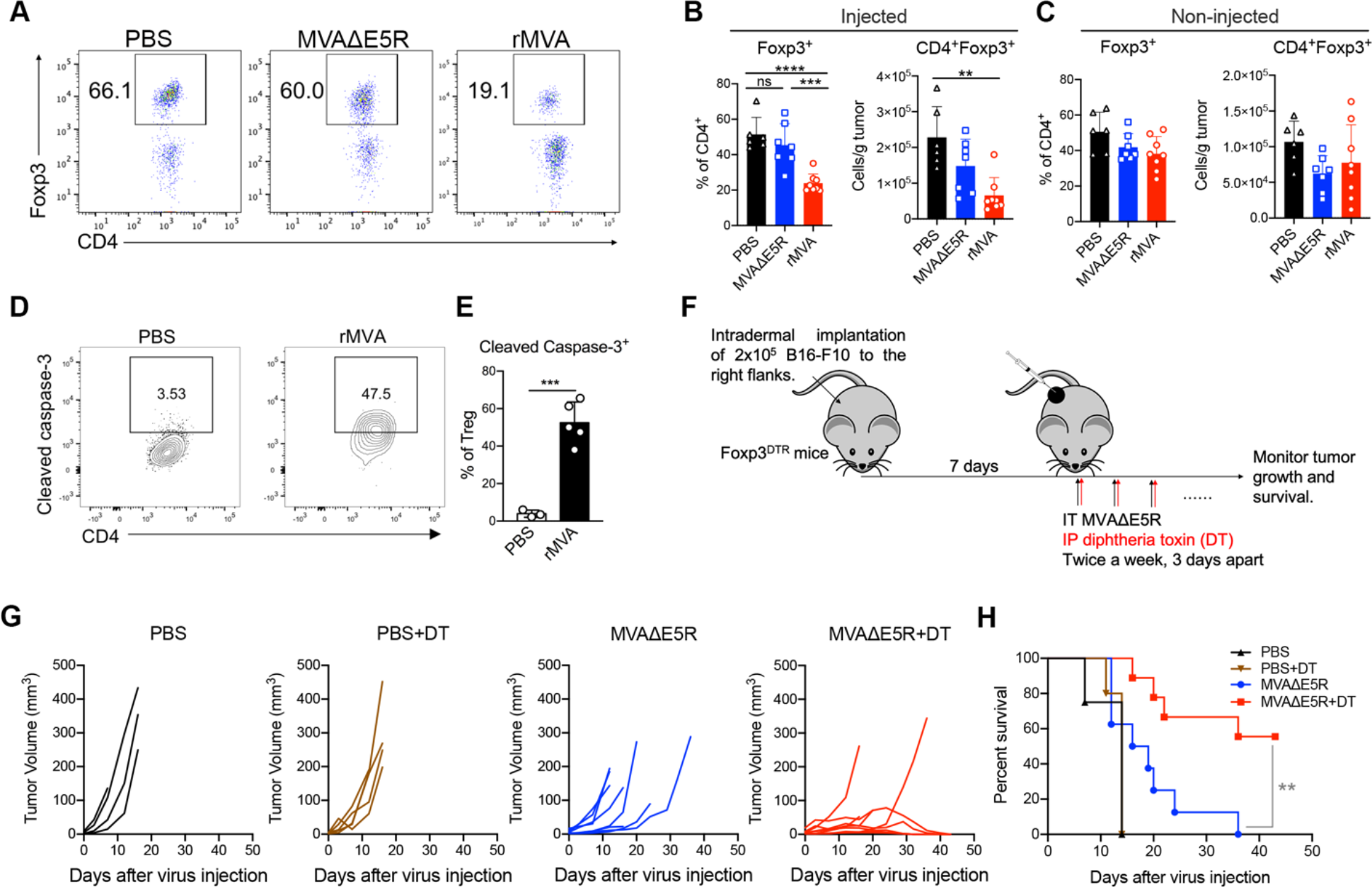
IT rMVA depletes OX40^hi^ Tregs in the injected tumors to promote anti-tumor therapy. (A) Representative flow cytometry plots of Foxp3^+^CD4^+^ cells in the injected tumors. Mice were treated as described in Fig. 3A. (B-C) Percentages and absolute number of Foxp3^+^CD4^+^ cells in the injected (B) and non-injected (C) tumors. Data are means ± SD (*n=6-8; **P < 0.01, ***P<0.001, ****P < 0.0001, t test).* (D-F) Mice were intradermally implanted with B16-F10 cells. Tumors were injected with rMVA or PBS as control after 7 days post implantation and harvested 2-days post injection. (D) Representative flow cytometry plots of cleaved caspase-3^+^ Tregs in the injected tumors. (E) ) Percentages of cleaved caspase-3 in tumor infiltrating-Tregs by flow cytometry. Data are means ± SD (*n=3-5; **P < 0.01, t test)*. (F) Schematic diagram of IT rMVA in the presence or absence of DT in a unilateral B16-F10 melanoma implantation model in Foxp3^DTR^ mice. (G) Tumor growth curves of mice treated with rMVA or PBS. (H) Kaplan-Meier survival curves of mice treated with rMVA or PBS (*n=5-10; **P < 0.01, Mantel-Cox test*).

To determine whether Tregs play a negative role in recombinant MVA-based virotherapy, we implanted B16-F10 cells intradermally into Foxp3-DTR mice. After the tumors were established, we treated the tumors with IT MVAΔE5R, which does not significantly reduce Tregs (**Figure 4F**). Although intraperitoneal administration of diphtheria toxin (DT) alone did not affect tumor growth or survival (**Figure 4G and 4H**), DT plus IT MVAΔE5R injection significantly improved therapeutic efficacy compared with IT MVAΔE5R alone (**Figure 4G and 4H**). These results suggest that Tregs play an inhibitory role in viral-based immunotherapy and intratumoral depletion of Tregs by rMVA is an important mechanism for potentiating antitumor immunity.

### IT rMVA preferentially depletes OX40^hi^ Tregs via OX40L-OX40 interaction and IFNAR signaling

We hypothesized that OX40L expressed by rMVA-infected myeloid and tumor cells might be important in mediating the reduction of OX40^hi^ Tregs. We first compared the surface expression of OX40 in various T cell populations within the tumor microenvironment. The mean percentages of OX40^hi^ Tregs among CD4^+^ Tregs were 51% compared with 5.6% of OX40^hi^ conventional T (Tcov) cells and 1% of OX40^hi^ CD8^+^ T cells (**Figure 5A).** The mean fluorescence intensity (MFI) of OX40 was much higher in CD4^+^ Tregs than those in Tcov and CD8^+^ T cells **(Figure 5B and 5C)**. The high expression of OX40 was unique to tumor-infiltrating Tregs, because OX40 expression levels in Tregs from spleens or lymph nodes (LNs) were much lower than those from tumors (**Figure S3A**). Next, we compared the tumor-infiltrating OX40^hi^ Treg population with or without IT rMVA treatment. IT rMVA treatment preferentially reduced the percentages of OX40^hi^ Tregs out of total Tregs, as well as the absolute numbers of OX40^hi^ Tregs per gram of tumors in the injected tumors (**Figure 5D and 5E**). In OX40-deficient mice, however, IT rMVA did not result in Treg reduction in the injected tumors (**Figure 5F**). Finally, we evaluated OX40L expression in both tumor cells and myeloid cells 2 days after IT rMVA. OX40L was detected on B16-F10, tumor-infiltrating macrophages, CD103^+^ DCs, CD11b^+^ DCs, neutrophils, and monocytes (**Figure S3B and S3C**). These results indicate that IT rMVA results in the OX40L expression in a variety of cell types including tumor and tumor-infiltrating myeloid cells and reduces tumor-infiltrating OX40^hi^ Tregs likely via OX40-OX40L interaction.

**Figure 5.**
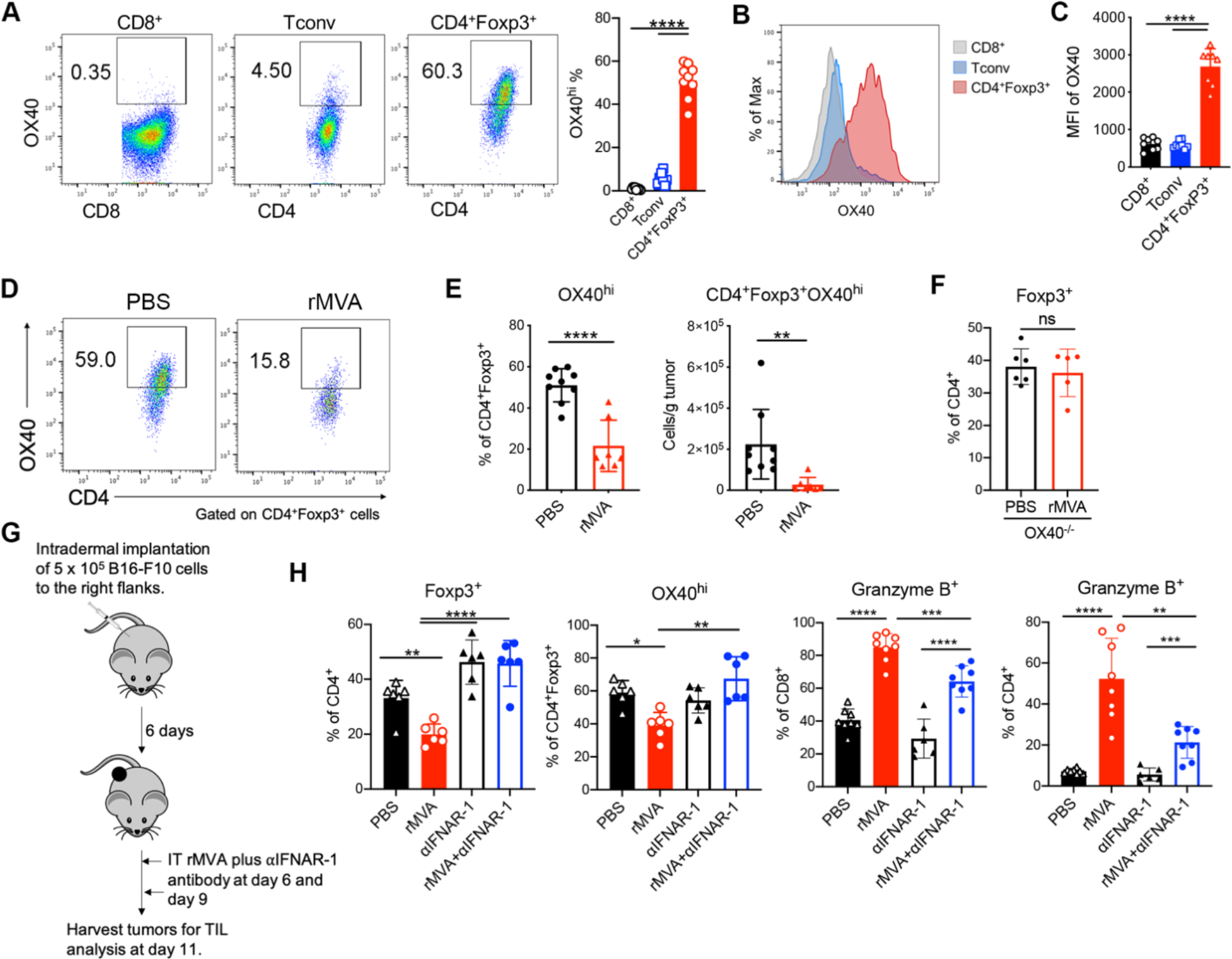
IT rMVA preferentially depletes OX40^hi^ Tregs in the injected tumors in a type I IFN signaling dependent manner. (A) Representative flow cytometry plots of OX40 expression on tumor infiltrating CD8^+^, Tconv, and CD4^+^Foxp3^+^ T cells in tumors 15 days after implantation. Mice were treated as described in Fig. 3A. (B-C) Representative flow cytometry plots and statistical analysis of mean fluorescence intensity of OX40 on tumor-infiltrating CD8^+^, Tconv, and CD4^+^Foxp3^+^ T cells. Data are means ± SD in (C) (*n=6∼8; ****P < 0.0001, t test)*. (D) Representative flow cytometry plots of OX40^hi^CD4^+^Foxp3^+^ in the injected tumors. Mice were treated as described in Fig. 3A. (E) Percentages and absolute number of OX40^hi^CD4^+^Foxp3^+^ T cells in the injected tumors. Data are means ± SD (*n=7 or 9; **P<0.01, ****P < 0.0001, t test)*. (F) Percentages of CD4^+^Foxp3^+^ T cells in the injected tumors from WT and OX40^-/-^ mice. Mice were treated as described in Fig. 3A. Data are means ± SD (*n=5 or 6; t test)*. (G) Schematic diagram of IT rMVA in the presence or absence of IT *α*IFNAR-1 antibody in a unilateral B16-F10 melanoma implantation model. (H) Percentages of CD4^+^Foxp3^+^ and OX40^hi^CD4^+^Foxp3^+^ T cells in the injected tumors. Data are means ± SD (*n=6; *P<0.05, **P<0.01, ****P < 0.0001, t test)*. (I) Percentages of CD8^+^Granzyme B^+^ and CD4^+^Granzyme B^+^ T cells in the injected tumors. Data are means ± SD (*n=6; **P<0.01, ***P<0.001, ****P < 0.0001, t test)*.

To evaluate whether the IFNAR signaling pathway is involved in rMVA-mediated Treg reduction, we co-administered anti-IFNAR-1 antibody with rMVA into implanted B16-F10 melanoma twice three days apart. Tumors were harvested two days post-second injection (**Figure 5G**). Whereas IT rMVA reduced the percentages of OX40^hi^ Treg out of Tregs as well as the percentages of Tregs out of CD4^+^ T cells, co-administration of anti-IFNAR-1 antibody with rMVA reversed the reduction (**Figure 5H**). In addition, co-administration of anti-IFNAR-1 with rMVA resulted in lower percentages of Granzyme B^+^ CD8^+^ and CD4^+^ T cells compared with IT rMVA alone (**Figure 5H**), which is consistent with the role of type I IFN in promoting CD8^+^ and CD4^+^ T cell activation. Taken together, our results provide strong evidence that IT rMVA results in the depletion of OX40^hi^ Tregs in the injected tumors via OX40L-OX40 interaction, and this process is facilitated by type I IFN induced by rMVA infection in the injected tumors.

### Tumor-infiltrating OX40^hi^ Tregs and OX40^lo^ Tregs have distinctive transcriptomic features

We observed that in the murine B16-F10 melanoma model, the percentage of OX40^hi^ Tregs in total Tregs positively correlated with tumor mass, suggesting that OX40^hi^ Tregs may represent an immunosuppressive cell population during tumor progression (**Figure 6A**). To determine the functional differences between OX40^hi^, OX40^lo^, and OX40^-/-^Tregs, we intradermally implanted B16-F10 cells into Foxp3^gfp^ and OX40^-/-^Foxp3^gfp^ mice in a C57BL/6J background and FACS-sorted OX40^hi^ and OX40^lo^ intratumoral Tregs from Foxp3^gfp^ mice and OX40^-/-^ Tregs from OX40^-/-^ Foxp3^gfp^ mice and compared their suppression function *in vitro*. Flow cytometry analysis showed that OX40^hi^ Tregs isolated from tumors suppressed Tconv proliferation more strongly compared with tumor-infiltrating OX40^lo^ Tregs or OX40^-/-^ Tregs *in vitro* (**Figure 6B and 6C**). We also isolated splenic Tregs from tumor-bearing Foxp3^gfp^ mice and found that the suppressive activities of these cells were similar to those of OX40^lo^ Tregs or OX40^-/-^ Tregs isolated from tumors (**Figure 6B and 6C**).

**Figure 6.**
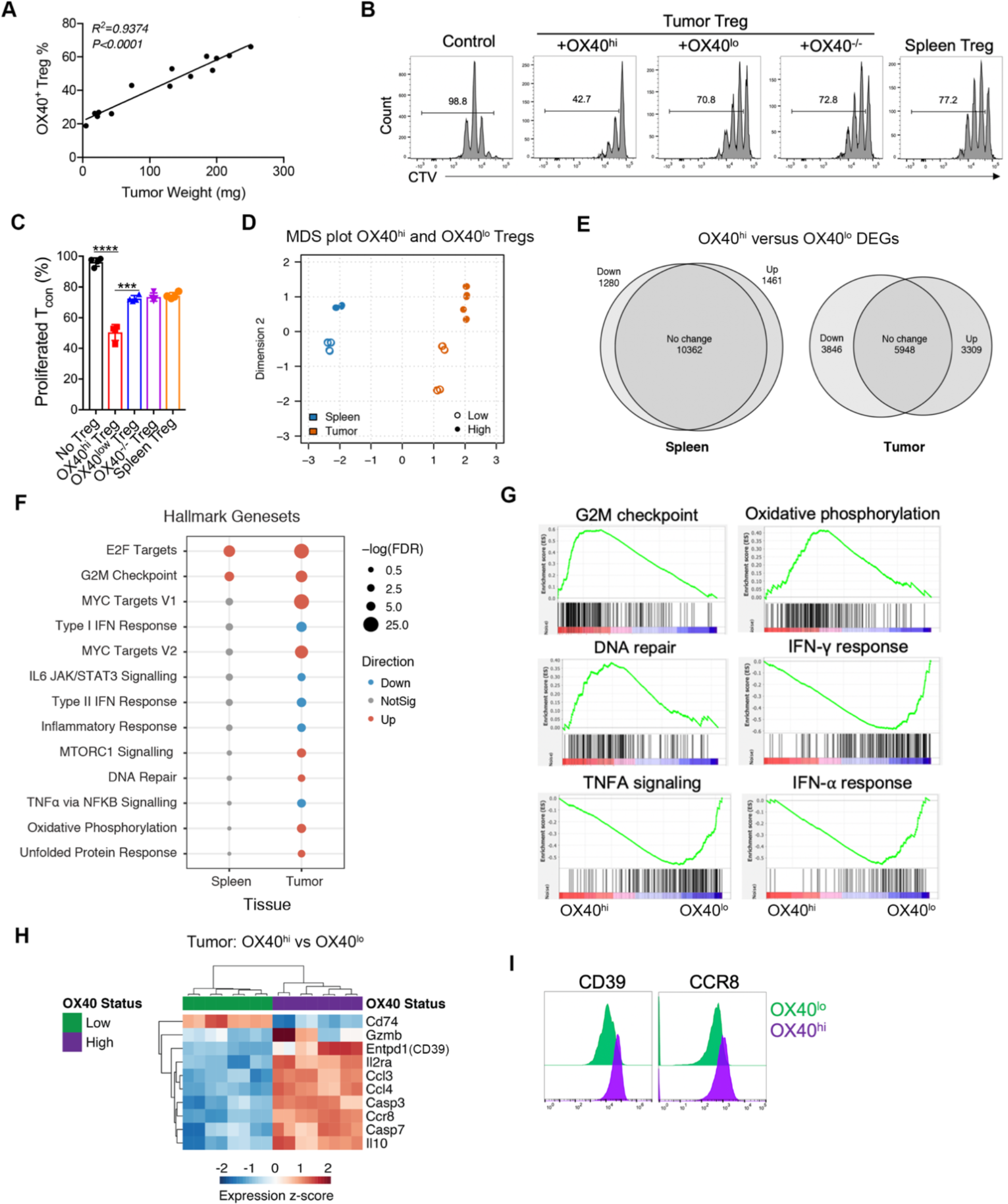
Tumor-infiltrating OX40^hi^ Tregs and OX40^low^ Tregs have distinctive transcriptomic features. (A) Correlation of the percentages of OX40^hi^ Tregs in the tumors with tumor weight. Mice were implanted with B16-F10 tumors intradermally. Tumors with different sizes were analyzed for OX40 expression on tumor-infiltrating Tregs. (B-C) Representative flow cytometry plots (B) and percentage of Tcon proliferation (C) as measured by CTV dye dilution co-cultured with tumor OX40^hi^, OX40^low^, OX40^-/-^ Tregs or spleen Tregs. Data are means ± SD in (C) (*\*\*\*P < 0.001; ****P < 0.0001, t test)*. (D) Multidimensional scaling (MDS) plot of RNA-seq results of OX40^hi^ and OX40^low^ Tregs from B16-F10 tumors. (E) Venn diagram of the relationship of differential gene expression (DEGs) of OX40^hi^ Tregs vs. OX40^lo^ Tregs isolated from spleens and tumors. (F) Gene set analysis of upregulated or downregulated genes comparing OX40^hi^ Tregs vs. OX40^lo^ Tregs isolated from spleens and tumors. (G) Gene Set Enrichment Analysis (GSEA) of upregulated or downregulated genes comparing transcriptomes of OX40^hi^ Tregs vs. OX40^lo^ Tregs isolated from tumors. (H) Heatmap of selected genes upregulated or downregulated genes in OX40^hi^ Tregs vs. OX40^lo^ Tregs isolated from tumors. (I) Representative FACS plot showing the expressions of CD39 and CCR8 on OX40^hi^ Tregs and OX40^lo^ Tregs isolated from tumors.

To explore the transcriptomic differences between OX40^hi^ and OX40^low^ Tregs isolated from tumors and spleens, RNA-seq analysis was performed on sorted Treg populations from Foxp3-GFP mice as described above. Multidimensional scaling (MDS) plot showed that OX40^hi^ and OX40^lo^ Tregs from either spleens or tumors were segregated into distinct populations (**Figure 6D**). Whereas splenic OX40^hi^ and OX40^lo^ Tregs were similar to each other at the transcriptomic level, intratumoral OX40^hi^ and OX40^lo^ Tregs diverged from each other with 3309 upregulated genes and 3846 down-regulated genes (**Figure 6E**). Pathway analysis showed that the main transcriptomic differences between splenic OX40^hi^ and OX40^lo^ Tregs were related to E2F targets and G2M check-points, suggesting that OX40^hi^ Tregs are more proliferative compared with OX40^lo^ Tregs in the spleens (**Figure 6F and 6G**). The upregulated genes in OX40^hi^ Tregs compared with OX40^lo^ Tregs from tumors belonged to the following pathways, including cell proliferation, DNA repair, oxidative phosphorylation, glycolysis, and the unfolded protein response (**Figure 6F and 6G**). Compared with OX40^lo^ Tregs in the tumors, OX40^hi^ Tregs had lower expression of genes that belong to type I and II IFN response and inflammation (**Figure 6F and 6G**). OX40^hi^ Tregs had higher expression of *Casp3* and *Casp7*, suggesting that they might be more prone to apoptosis (**Figure 6H)**. In addition, OX40^hi^ Tregs had higher *Il2ra* expression compared with OX40^lo^ Tregs, which may explain their stronger suppressive capacity through competition for IL2 in the tumor micro-environment (**Figure 6H**). OX40^hi^ Tregs also had high expression of *Ccl3* and *Ccl4* chemokines (**Figure 6H**), which have been implicated in chemoattraction of CCR5^+^ CD4^+^ and CD8^+^ T cells and immune suppression (Patterson et al., 2016). FACS analyses confirmed higher *CD39* and *Ccr8* expression in OX40^hi^ tumor-infiltrating Tregs compared with OX40^lo^ tumor-infiltrating Tregs (**Figure 6I**).

Comparison of OX40^hi^ Tregs in tumors vs. spleens revealed striking differences in these two populations. OX40^hi^ Tregs in tumors have higher expression of chemokine receptors including *Ccr2, Ccr5, Ccr8, Cxcr3, and Cxcr6* as well as chemokines including *Ccl1, Ccl2, Ccl3, Ccl4, Ccl5, Ccl7, Ccl8, Ccl12, Ccl17, Ccl22, Cxcl2, and Cxcl9*, which suggest that they are migratory in response to chemokine cues in the developing tumors (**Figure S4A**). OX40^hi^ Tregs in tumors also express higher levels of *Il10, Ctla4, Tnfrsf18 (GITR)*, thus correlating with a more potent immunosuppressive function (**Figure S4B**). Furthermore, OX40^hi^ Tregs in tumors express higher levels of genes involved in glycolysis and oxidative phosphorylation (**Figure S4C and S4D**).

### CD8^+^ T cells are required for the antitumor effects induced by IT rMVA

To determine which cell populations are essential for tumor eradication by IT rMVA, we performed an antibody depletion experiment using anti-CD8 and/or anti-CD4 antibodies during rMVA treatment. The depleting antibodies were first given intraperitoneally two days before IT rMVA and then were given at the same time when mice were treated with IT rMVA (**Figure 7A**). Depletion of CD8^+^ cells abrogated the therapeutic effect of rMVA (**Figure 7B and 7C**). Although CD4^+^ T cell depletion resulted in a better response to rMVA treatment initially, however, the antitumor response did not persist in the CD4^+^ T cell-depleted mice after IT rMVA treatment ended on day 42. 60% of mice died due to the recurrence of tumors. Mice with both CD4^+^ and CD8^+^ T cell-depleted behaved similarly to those with just CD8^+^ T cell depletion (**Figure 7B and 7C**). These results suggested that CD8^+^ T cells are required for tumor eradication in IT rMVA therapy, while CD4^+^ T cells may be important for facilitating the generation of anti-tumor memory responses.

**Figure 7.**
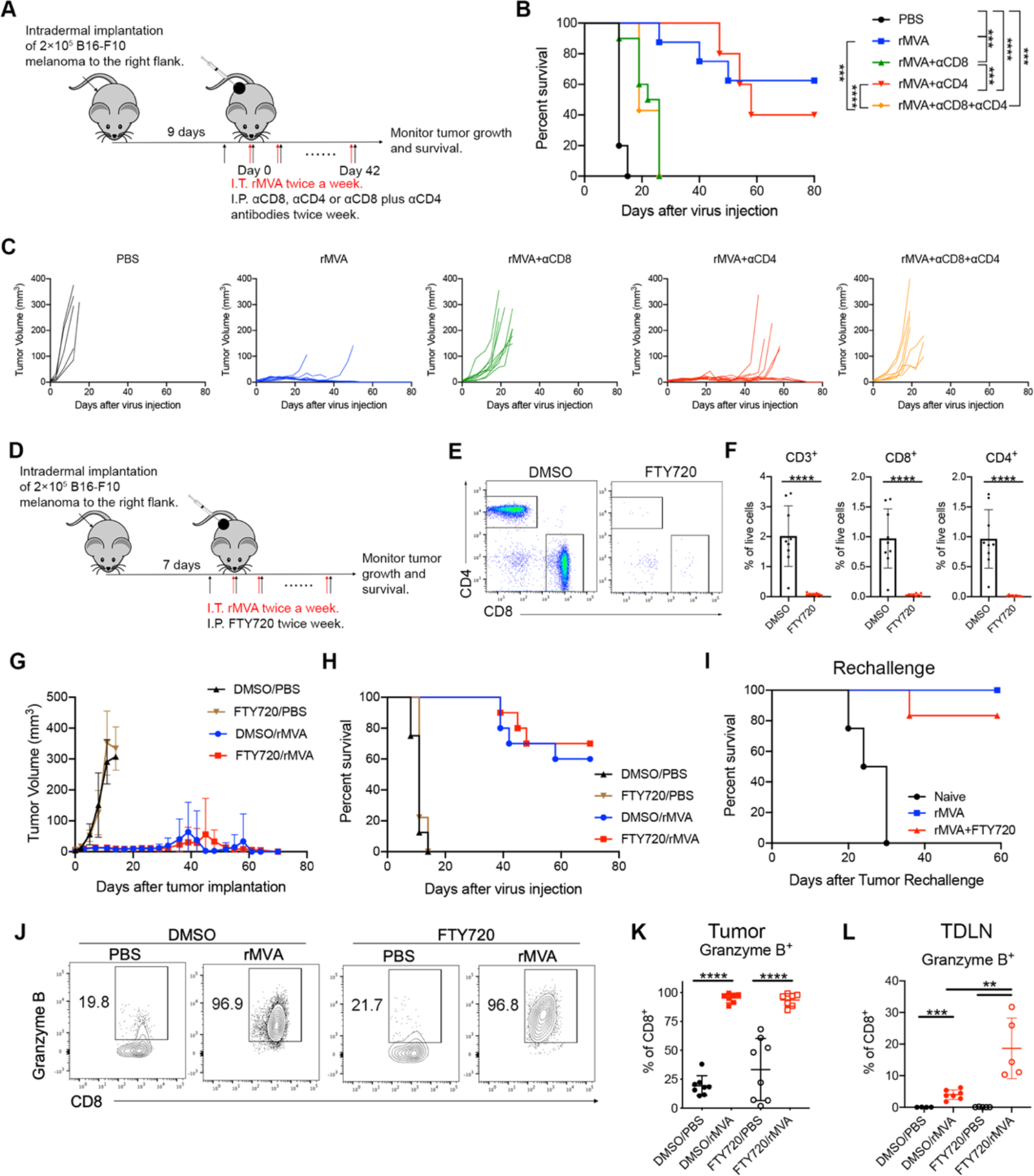
Local activation of CD8^+^ T cells is sufficient for rMVA-mediated tumor eradication. (A) Schematic diagram of IT rMVA in the presence or absence of depleting antibodies for CD8 and/or CD4 in a unilateral B16-F10 melanoma implantation model. (B) Kaplan-Meier survival curves (*n=5∼10; ***P < 0.001, ****P < 0.0001, Mantel-Cox test*). (C) Tumor growth curves. (D) Schematic diagram of IT rMVA in the presence or absence of FTY720 in a unilateral B16-F10 melanoma implantation model. (E) Representative flow cytometry plot of CD8**^+^** and CD4**^+^** T cells in the PBL of mice treated with FTY720. (F) Percentages of CD8**^+^** and CD4**^+^** T cells in the PBL of mice treated with FTY720. Data are means ± SD (*n=10; ****P < 0.0001, t test)*. (G) Tumor growth curves. (H) Kaplan-Meier survival curves (*n=10*). (I) Kaplan-Meier survival curves of survived mice from (H) or naïve mice challenged with 1x10^5^ B16-F10 cells at the contralateral side (*n=5 or 6*). (J) Representative flow cytometry plots of Granzyme B^+^CD8^+^ T cells in the injected tumors. (K) Percentages of Granzyme B^+^CD8^+^ T cells in the injected tumors. Data are means ± SD (*n=8; ****P < 0.0001, t test)*. (L) Percentages of Granzyme B^+^CD8^+^ T cells in the tumor-draining lymph nodes. Data are means ± SD (*n=6∼8; **P < 0.01, ***P < 0.001, t test)*.

### IT rMVA activates pre-existing CD8^+^ T cells in the tumors, which is sufficient for the eradication of injected tumors

To determine whether local T cell activation or T cell recruitment from tumor-draining lymph nodes (TDLNs) is important for rMVA-induced antitumor effects in the injected tumors, we administered FTY720 (fingolimod), a modulator of the sphingosine-1-phosphate receptor, which blocks T cell egress from lymphoid organs. FTY720 was given intraperitoneally one day before IT rMVA and was later given to the mice twice a week on the days when they were treated with IT rMVA (**Figure 7D**). FACS analysis confirmed that FTY720 treatment depletes CD8^+^ and CD4^+^ T cells in the blood (**Figure 7E and 7F**). We observed that FTY720 treatment alone did not affect tumor growth. Co-administration of FTY720 with IT rMVA treatment did not diminish the antitumor effect of IT rMVA (**Figure 7G and 7H**). When the suriving mice previously treated with IT rMVA alone or with IT rMVA plus FTY720 were challenged with a lethal dose of B16-F10 on the contralateral side at 7 weeks post tumor eradication, 80% of the mice in the rMVA plus FTY720 group and 100% of the mice in the rMVA alone group were able to reject tumor challenge (**Figure 7I**). No FTY720 was administered during the challenge phase. We analyzed T cells from both the injected tumors and TDLNs with or without FTY720 treatment, and found that Granzyme B^+^ CD8^+^ T cells were increased after rMVA treatment with or without FTY720, indicating that IT rMVA directly activates preexisting antitumor CD8^+^ T cells in the tumors without lymph node involvement (**Figure 7J and 7K**). In the FTY720- treated group, the percentages of Granzyme B^+^CD8^+^ T cells in the TDLNs were higher than that of the DMSO control group with rMVA injection (**Figure 7L**), indicating that the newly primed CD8^+^ T cells were trapped in the TDLNs. These results indicate that IT rMVA-induced local activation of anti-tumor T cells is sufficient for the eradication of injected tumors.

### Combination of IT delivery of rMVA with systemic administration of anti-PD-L1 antibody provides systemic antitumor therapeutic effects

B16-F10 tumors respond poorly to immune checkpoint blockade (ICB) therapy. To test whether the combination of systemic delivery of ICB antibody and IT rMVA therapy can overcome the resistance to ICB therapy, we used a bilateral B16-F10 implantation model and compared the antitumor efficacy of with IT rMVA alone vs. IT rMVA plus intraperitoneal (IP) delivery of anti-PD-L1 antibody (**Figure 8A**). IT rMVA alone eradicated 9 out of 10 injected tumors and delayed the growth of non-injected tumors. However, 90% of mice died eventually due to the growth of non-injected tumors (**Figure 8B and 8C**). By contrast, the combination of IP anti-PD-L1 antibody and IT rMVA significantly improved the antitumor therapeutic efficacy (**Figure 8B and 8C**). 80% of mice in the combination group rejected non-injected tumors and survived. These results demonstrated that the combination of systemic delivery of anti-PD-L1 antibody and IT rMVA generates synergistic systemic antitumor therapeutic effects, leading to the eradication of both injected and non-injected tumors.

**Figure 8.**
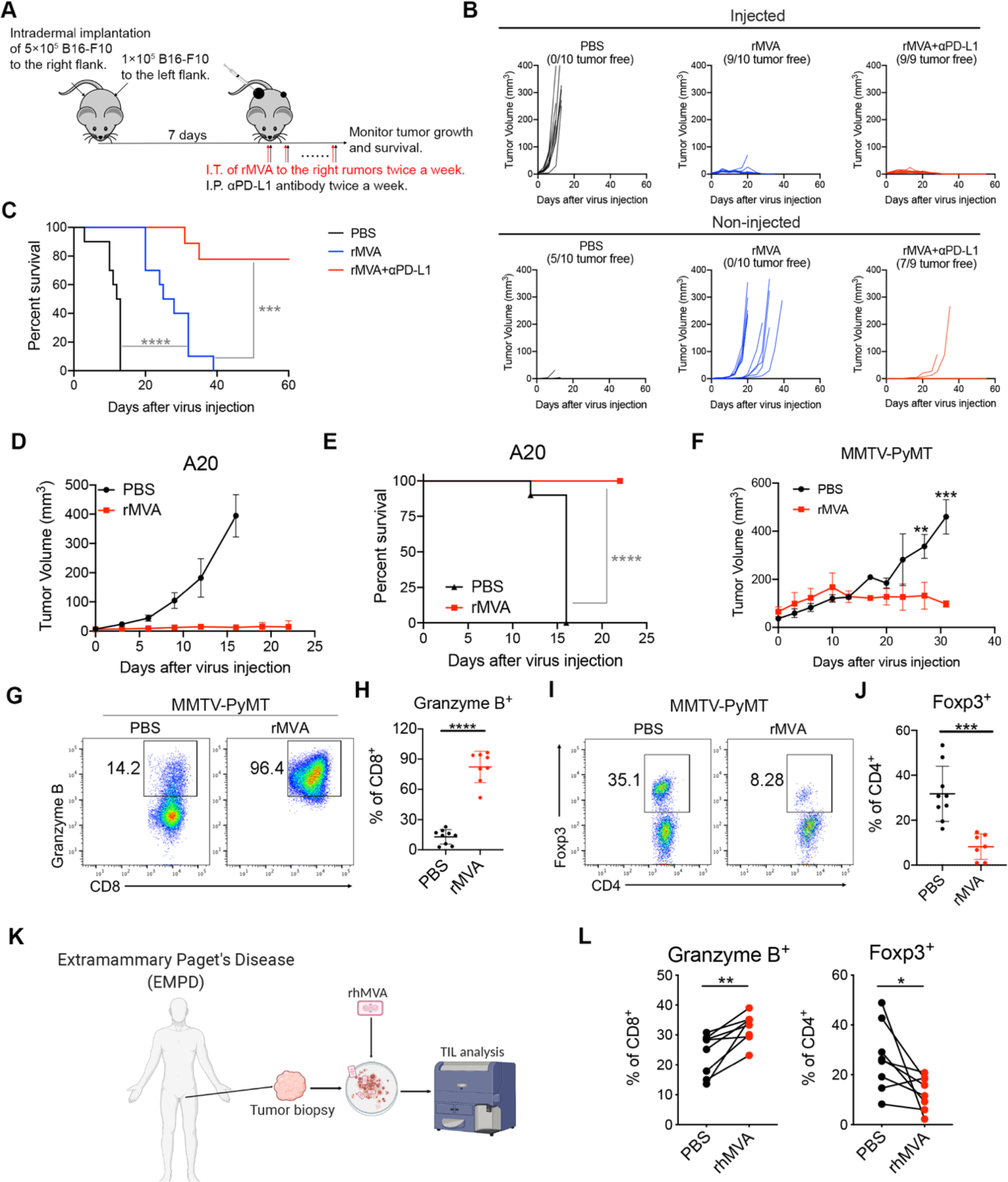
IT rMVA elicits strong antitumor immunity in multiple murine tumor models. (A) Schematic diagram of IT rMVA in combination with IP *α*PD-L1 antibody in a bilateral B16-F10 melanoma implantation model. (B) Tumor growth curves. (C) Kaplan-Meier survival curves (*n=9 or 10; **** P < 0.0001, Mantel-Cox test*). (D) Tumor growth curves of mice treated with IT rMVA in an A20 B cell lymphoma implantation model. (E) Kaplan-Meier survival curves (*n=10; ****P < 0.0001, Mantel-Cox test*). (F) Tumor growth curves in the PyMT-MMTV breast tumor model. Data are means ± SD (*n=5; **P < 0.01, ***P < 0.001, t test)*. (G) Representative flow cytometry plots of Granzyme B^+^CD8^+^ T cells in the IT rMVA-treated tumors from MMTV-PyMT mice. (H) Percentages of Granzyme B^+^ CD8^+^ T cells in the injected tumors. Data are means ± SD (*n=8; ****P < 0.0001, t test)*. (I) Representative flow cytometry plots of Foxp3^+^CD4^+^ T cells in the injected tumors. (J) Percentages of Foxp3^+^CD4^+^ T cells in the injected tumors. Data are means ± SD (*n=8; ***P < 0.001, t test)*. (K) Schematic diagram of ex vivo infection of human EMPD tumors with rhMVA. (L) Percentages of Granzyme B^+^CD8^+^ T cells and Foxp3^+^CD4^+^ T cells in the rhMVA-treated and non-treated tumor tissues. Data are means ± SD (*n=7; *P < 0.05, **P < 0.01, t test)*.

### IT rMVA is effective in generating antitumor T cell responses and controlling tumor growth in murine A20 B cell lymphoma and triple-negative breast cancer models

In addition to murine B16-F10 melanoma, we evaluated the therapeutic efficacy of IT rMVA in other murine tumor models. IT rMVA efficiently eradicated A20 B cell lymphoma tumors and resulted in 100% survival (**Figure 8D and 8E**). MMTV-PyMT is a transgenic mouse strain that develops multiple tumors in the mammary fat pads spontaneously, commonly used as a triple-negative breast tumor model. In the MMTV-PyMT mice, IT injection of rMVA resulted in delayed tumor growth compared with the PBS control group (**Figure 8F**). Similar to what we observed in the B16-F10 murine melanoma model, IT rMVA activated CD8^+^ T cells in the tumors and reduced Tregs (**Figure 8G-8I**).

### Clinical candidate rhMVA (MVAΔE5R-hFlt3L-hOX40L) induces innate immunity and promotes maturation of human monocyte-derived DCs (moDCs)

For clinical applications, we generated an rhMVA expressing human Flt3L and human OX40L and with the deletion of E5R gene (**Figure S5A**). hFlt3l and hOX40L are membrane-bound ligands that were expressed on the surface of murine B16-F10 cells and human melanoma cell line, SK-MEL-28, after infection with rhMVA *in vitro* (**Figure S5B**). rhMVA induced higher levels of *ifnb* gene expression as well as *ccl4, ccl5, cxcl10, il1b, il6*, and *tnf* in moDCs compared with MVA (**Figure S5C**). rhMVA infection of moDCs induced the expression of CD86 on the cell surface, which is indicative of DC maturation (**Figure S5D**).

To test whether *ex vivo* infection of human tumor samples with rhMVA could induce phenotypic changes of tumor-infiltrating lymphocytes (TILs), we obtained skin biopsy samples from patients with extramammary paget’s disease (EMPD), infected the processed tissues with rhMVA, and analyzed TILs 24 h later (**Figure 8K**). We found that rhMVA-infected samples exhibited upregulation of granzyme B on CD8^+^ T cells as well as reduction of Tregs compared with the paired control samples (**Figure 8L**). These results are consistent with what we observed in various murine tumors treated with rMVA *in vivo*, further supporting rhMVA as a potential clinical candidate for the treatment of various human cancers.

## Discussion

Preclinical and clinical studies have shown that viral-based immunotherapeutics can alter immunosuppressive tumor microenvironment to enhance antitumor effects and overcome resistance to immune checkpoint blockade antibody therapy. Our previous study using heat-inactivated modified vaccinia virus Ankara (heat-iMVA) demonstrated that induction of innate immunity via the STING-dependent pathway in the tumor microenvironment is important for the generation of systemic antitumor immunity (Dai et al., 2017), which is also dependent on CD8^+^ T cells and Batf3dependent CD103^+^/CD8*α*^+^ DCs. Based on this concept, we engineered a recombinant MVA (rMVA) with deletion of the vaccinia E5R gene, which encodes an inhibitor of cGAS, and with the expression of two membrane-bound transgenes, human Flt3L, and murine OX40L. Here we show that rMVA activates innate immunity via the cGAS/STING pathway and the IFNAR positive feedback loop, and also reduces OX40^hi^ regulatory T cells via OX40L-OX40 interaction in the injected tumors.

Poxviruses are large cytoplasmic DNA viruses. DNAs from the parental viral genome and the replicated progeny viral genome are potent stimuli for activating the cytosolic DNA-sensing pathway mediated by cGAS/STING. The vaccinia E5R gene is highly conserved among the orthopoxvirus family which includes variola virus, the causative agent of smallpox, and vaccinia virus, the laboratory strain that leads to the eradication of smallpox. MVAΔE5R infection of BMDCs induces higher levels of type I IFN compared with MVA, which is dependent on cGAS. Similarly, rMVA infection of BMDCs induces much higher levels of IFN compared with MVA. rMVA infection also induces cGAS-dependent DC maturation. IT delivery of rMVA generates stronger local and systemic antitumor effects compared with MVA, or MVAΔE5R. Using cGAS, STING, STAT1, and STAT2-deficient mice, we demonstrated that the cGAS/STING-mediated cytosolic DNA-sensing and STAT1/STAT2-mediated type I IFN pathways are required for the generation of antitumor immunity by rMVA. In addition to the induction of innate immunity, IT delivery of rMVA reduces OX40^hi^ Foxp3^+^CD4^+^ regulatory T cells in the injected tumors via OX40L/OX40 interaction which is reversed in the presence of anti-IFNAR antibody. Furthermore, using FTY720 to block lymphocyte egress from lymphoid organs, we demonstrated that IT rMVA-induced local activation of tumor-specific T cells in the injected tumors is sufficient for its eradication.

To understand why the cGAS/STING pathway is important in rMVA-induced antitumor immunity, we first evaluated what cell populations in the tumor microenvironment are preferentially infected by the virus. Using MVAΔE5R-expressing the mCherry reporter under the vaccinia virus synthetic early/late promoter, we found that the myeloid cell populations including macrophages, monocytes, neutrophils, and dendritic cells are targeted by the virus, whereas lymphocytes are largely spared. RNA-seq analyses of tumor tissues isolated 24 h post IT rMVA revealed marked upregulation in the expression of *Ifnb*, inflammatory cytokines and chemokines, DC activation markers, IFN-stimulated genes, and genes involved in apoptosis in WT mice, but not in STING-deficient mice. On the contrary, the expression of genes involved in oxidative phosphorylation was reduced in rMVA-treated tumors in WT mice, but not in STING-deficient mice.

We next addressed how innate immune activation, for example, the cGAS/STING pathway and type I IFN signaling in the tumor microenvironment, affects the activation status of tumor-infiltrating T cells. Using cGAS, STING, or STAT2-deficient mice or intratumoral delivery of anti-IFNAR antibody, we found that the activation of intratumoral CD8^+^ and CD4^+^ T cells depends on the generation of type I IFN and IFN signaling in the tumor microenvironment. This is consistent with a published report that type I IFN signaling drives antigen-independent expression of granzyme B on memory CD8^+^ T cells in a respiratory viral infection model (Kohlmeier et al., 2010). While IT rMVA was ineffective in shrinking tumors when CD8^+^ T cells were depleted, CD4^+^ T cells depletion did not impede tumor control initially. However, tumors regrew when both the anti-CD4 antibody and rMVA treatment were discontinued. We interpret these results as the following: As anti-CD4 antibody removes both Tcov and Tregs from tumors and circulation, mice were able to control tumor growth with activated CD8^+^ T cells which function better in the absence of Tregs. However, they fail to develop antitumor memory CD8^+^ T cells in the absence of helper CD4^+^ T cells.

Our study demonstrates that IT rMVA preferentially reduces OX40^hi^ Tregs in injected tumors via the OX40L-OX40 interaction and this process is promoted by type I IFN in the tumor microenvironment. We found that OX40 is preferentially expressed by intratumoral Tregs and its expression correlates with tumor weight in murine melanoma models. OX40^hi^ Tregs isolated from tumors are more immunosuppressive compared with OX40^low^ Tregs. Comparison of the transcriptomes of OX40^hi^ and OX40^lo^ intratumoral Tregs revealed that OX40^hi^ Tregs are more proliferative, metabolically active, and immunosuppressive. Therefore, targeting intratumoral OX40^hi^ Tregs by rMVA expressing OX40L is a logical approach to deplete this cell population within the tumors but not in the periphery, and thereby improving the efficacy of immunotherapy without unwanted autoimmunity. We found that OX40^hi^ Tregs expressed higher levels of CCR8 compared with OX40^lo^ Tregs. CCR8^+^ Tregs have been reported to play important roles in immune suppression in mice and humans (Barsheshet et al., 2017; Coghill et al., 2013; Plitas et al., 2016). Targeting CCR8^+^ Tregs using an anti-CCR8 antibody showed therapeutic benefits for cancer treatment and cancer vaccines in preclinical models (Villarreal et al., 2018). In addition to CCR8^+^, OX40^hi^ Tregs also expressed higher levels of *Il2ra, Ctla4, Tnfrsf18, Il10*, and *Cd39*, which are consistent with their immunosuppressive functions (Chinen et al., 2016; Cohen et al., 2010; Leone et al., 2018; Maj et al., 2017; Wing et al., 2008; Zappasodi et al., 2019).

Our results also showed that intratumoral OX40^hi^ Tregs had marked upregulation of genes involved in oxidative phosphorylation compared with intratumoral OX40^lo^ Tregs or with OX40^hi^ Tregs in the spleens. Several recent studies highlight the potential of modulating Treg stability and functions through metabolic control (Field et al., 2020; He et al., 2017; Weinberg et al., 2019; Zappasodi et al., 2021). How IT rMVA alters metabolism of OX40^hi^ Tregs via OX40L-OX40 interaction and the role of IFNAR signaling on Tregs on metabolic alterations and apoptosis will be addressed in future studies.

FTY720 (fingolimod), a sphingosine 1-phosphate receptor (S1P-R) agonist, is a novel immunomodulatory agent that blocks lymphocyte egress from lymphoid organs, FDA-approved for the treatment of relapsing multiple sclerosis. We observed that FTY720 did not affect the antitumor effects of rMVA in the injected tumors, despite trapping CD8^+^ T cells in the TDLNs, supporting our hypothesis that local activation of CD8^+^ T cells in the injected tumors is sufficient to eliminate the injected tumors. We provided evidence that type I IFN induction by rMVA in the injected tumors facilitates the proliferation and activation of CD8^+^ T cells. In addition, IFN contributes to the depletion of OX40^hi^ Tregs via the OX40L-OX40 ligation on Tregs. Removal of immunosuppressive OX40^hi^ Tregs in the rMVA-infected tumors leads to the activation of CD8^+^ and CD4^+^ T cells.

Taken together, we propose the following working model for elucidating the mechanisms of action of rMVA (**Figure S6**). After intratumoral injection with rMVA, both tumors and tumor-infiltrating myeloid cells are infected by the virus, which leads to the expression of hFlt3L and mOX40L transgenes on the cell surface, as well as the induction of type I IFN and proinflammatory cytokines and chemokines via the cGAS/STING-dependent cytosolic DNA-sensing pathway from the resident and recruited tumor-infiltrating myeloid cells. Type I IFN plays an important role in activating DCs and tumor-infiltrating CD8^+^ and CD4^+^ T cells. In addition, OX40L expression on infected tumor and myeloid cells leads to the depletion of OX40^hi^ Tregs via OX40L-OX40 interaction, which further enhances the antitumor activities of CD8^+^ and CD4^+^ T cells. Therefore, rMVA engages both innate and adaptive immunity to generate local and systemic antitumor effects, which are amplified in the presence of immune checkpoint blockade antibodies.

OX40 modulating agents have been explored for enhancing antitumor effects through targeting Tregs. For example, the combination of anti-OX40 agonist antibody with cyclophosphamide triggers activation and apoptosis of intratumoral Tregs while causing Treg expansion in the TDLN and spleens (Hirschhorn-Cymerman et al., 2009). This combination also promotes tumor-killing activities of antigen-specific adoptively transferred CD4^+^ T cells (Hirschhorn-Cymerman et al., 2012). In addition, intratumoral delivery of low doses of anti-CTLA-4 and anti-OX40 antibodies together with TLR9 agonist CpG leads to Treg depletion at the injected site but not in the non-injected site and generates systemic antitumor immunity (Marabelle et al., 2013). Our RNA-seq results confirmed that OX40^hi^ intratumoral Tregs express higher levels of CTLA-4 as well as other immune suppressive markers compared with OX40^lo^ Tregs. Therefore, our approach of using a recombinant immune-activating virus to express OX40L represents a novel strategy to deplete OX40^hi^ immunosuppressive Tregs. Compared with anti-OX40 agonist antibody approach, our engineered virus expressing OX40L might be more specific in targeting OX40^hi^ Tregs within the tumor microenvironment. IFN-inducing ability of the virus also promotes Treg depletion. The combination of intratumoral delivery of rMVA expressing mOX40L with systemic delivery of anti-PD-L1 antibody generates synergistic antitumor effects. This is in contrast to two reports showing that concurrent administration of anti-PD1 antibody leads to reduced efficacy of anti-OX40 antibody due to apoptosis of activated T cells (Messenheimer et al., 2017; Shrimali et al., 2017).

## STAR Methods

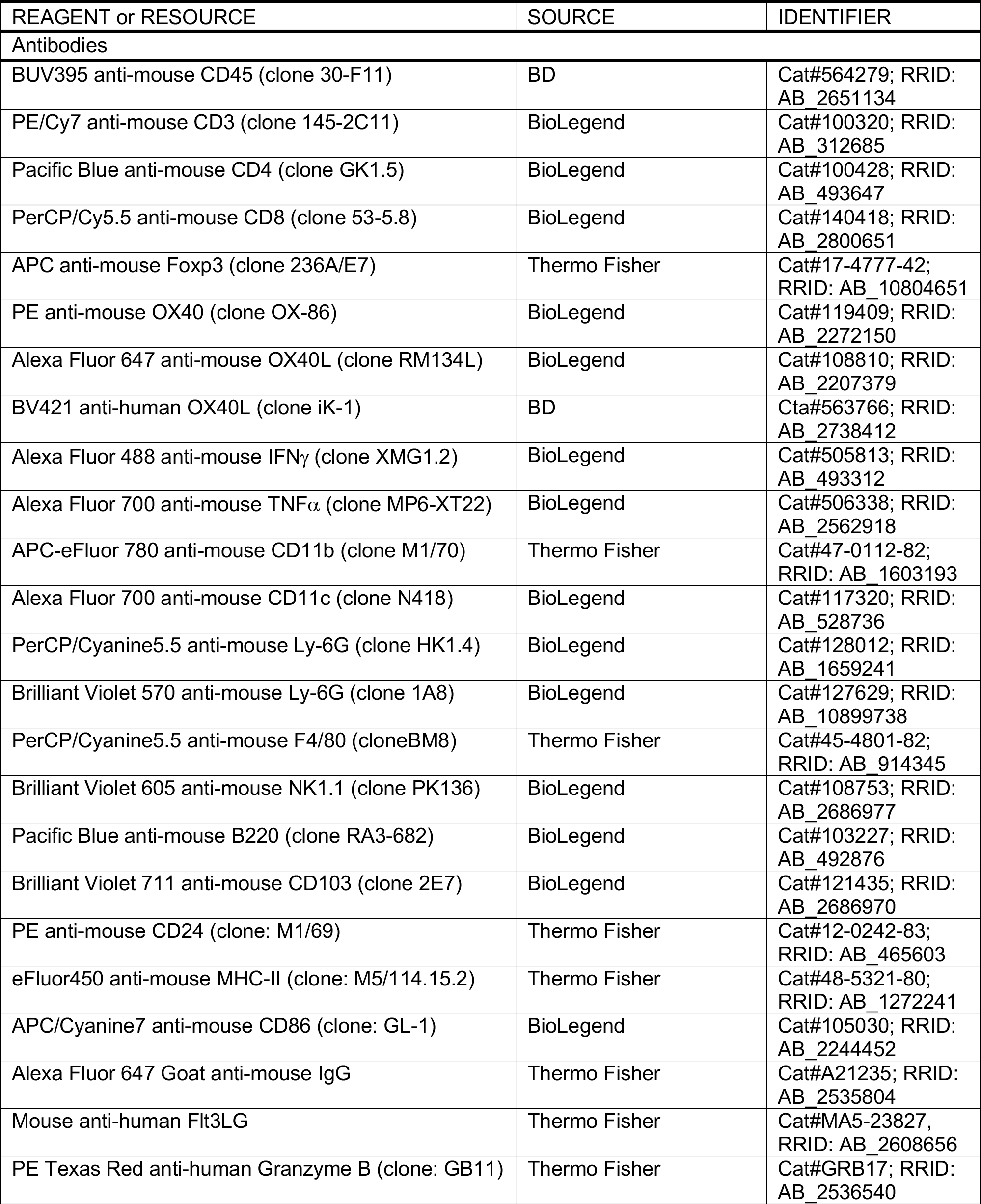
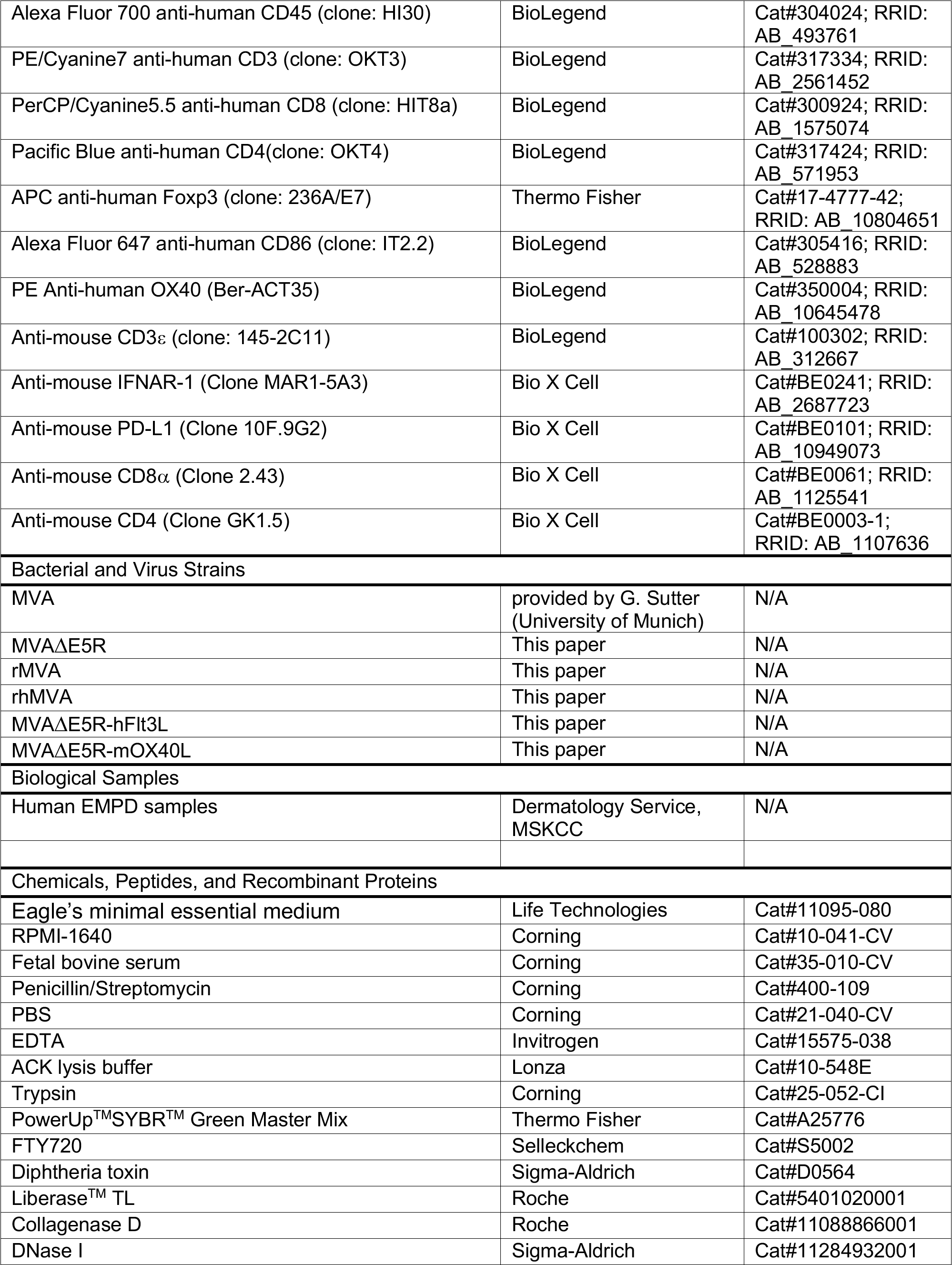
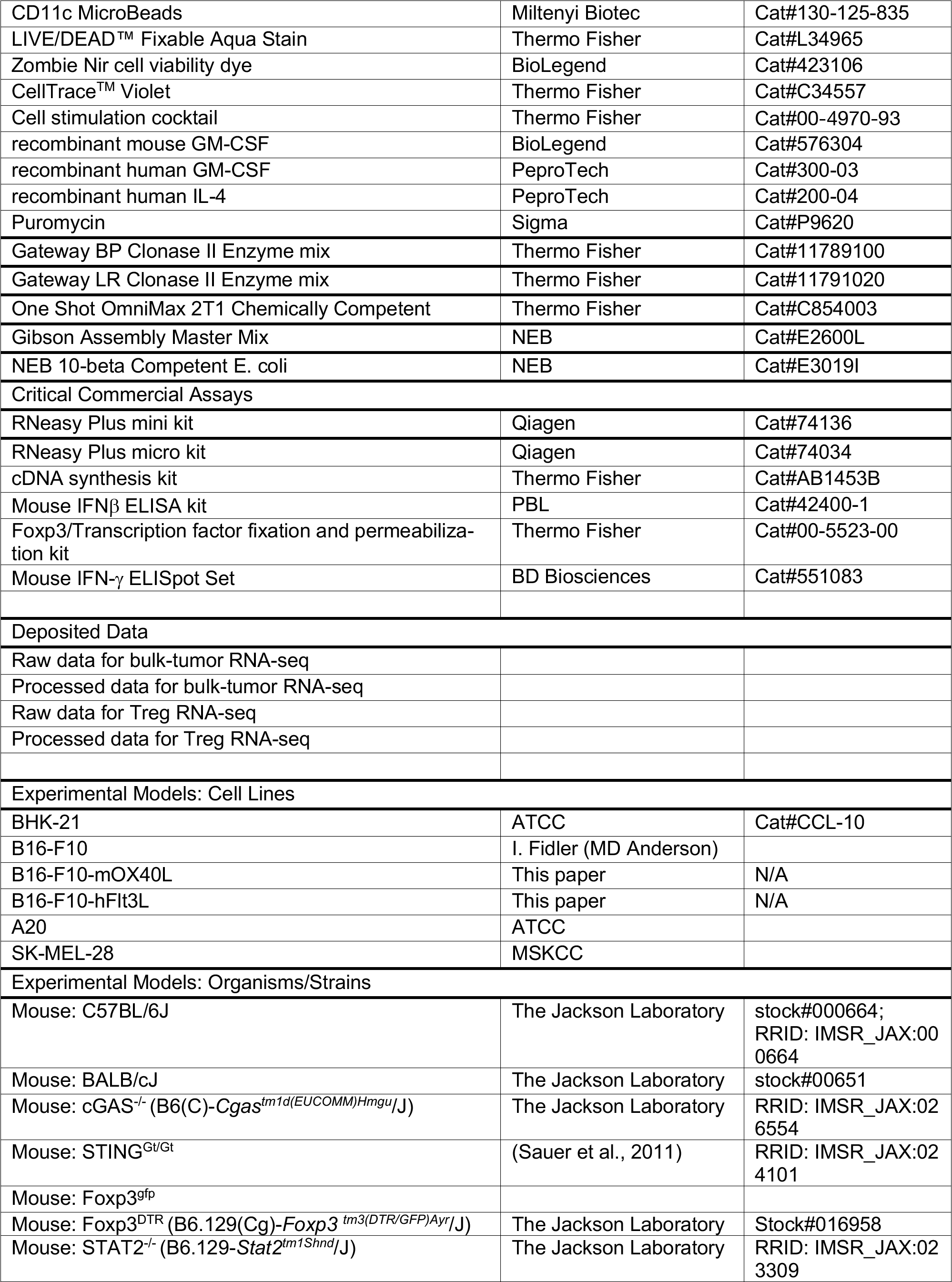
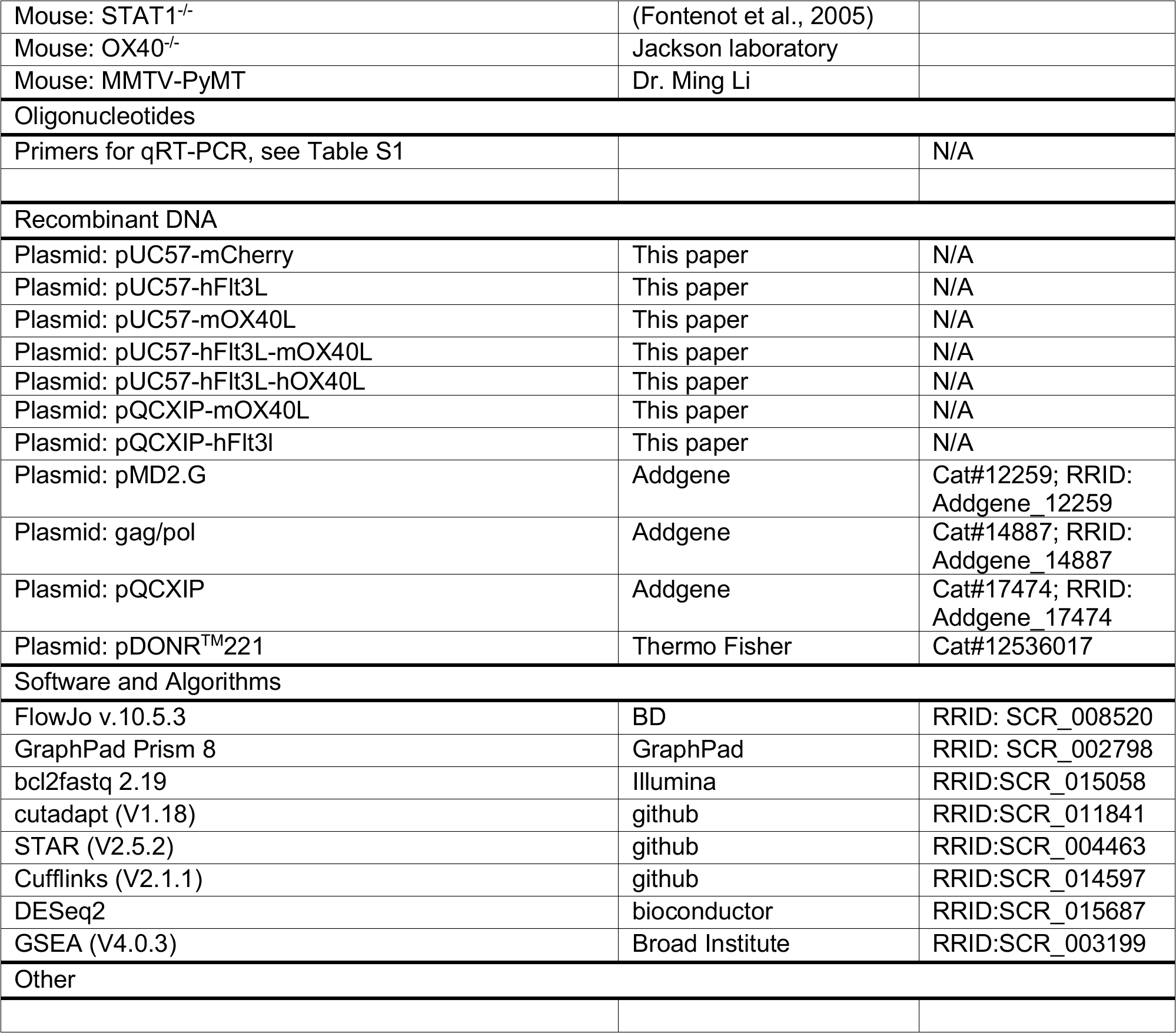
KEY RESOURCES TABLE

### Resource availability

#### Lead Contact

Further information and requests for resources and reagents used in this study should be directed to and will be fulfilled by the lead contact and the corresponding author, Liang Deng.

#### Materials availability

The materials used in this study are listed in the Key Resources Table. Materials generated in our laboratory are available upon request.

##### Data and code availability

The RNA-seq data reported in this study have been deposited in the Gene Expression Omnibus database (GEO) under the accession number GEO: ---. We analyzed the RNA-seq data ---

### Materials and Methods

#### Cell lines

BHK-21 (baby hamster kidney cell, ATCC CCL-10) cells were cultured in Eagle’s minimal essential medium containing 10% fetal bovine serum (FBS), 0.1 mM nonessential amino acids, penicillin, and streptomycin. The murine melanoma cell line B16-F10 was originally obtained from I. Fidler (MD Anderson Cancer Center). The A20 B cell lymphoma cell line were obtained from ATCC. Both B16-F10 and A20 were maintained in RPMI-1640 medium supplemented with 10% FBS, 0.05 mM 2-mercaptoethanol, penicillin, and streptomycin.

B16 cell line expressing murine OX40L (mOX40L) or human Flt3l (fFlt3l) were created by transduction into B16 cells with vesicular stomatitis virus (VSV) G protein-pseudotyped murine leukemia viruses (MLV) containing pQCXIP-mOX40L or pQCXIP-fFlt3l. Cells were selected and maintained in growth media including 2 μg/ml puromycin for selection of stably transduced cells.

#### Viruses

The MVA virus was provided by G. Sutter (University of Munich). MVA*Δ*E5R, MVA*Δ*E5R-hFlt3L, MVA*Δ*E5R-mOX40L, rMVA and rhMVA were generated by transfecting pUC57-based plasmids into BHK-21 cells that were infected with MVA at MOI 0.05. Recombinant viruses were purified after 4∼6 rounds of plaque selection based on the fluorescence marker. Viruses were propagated in BHK-21 cells and purified through a 36% sucrose cushion.

#### Mice

Female C57BL/6J mice and BALB/cJ between 6 and 10 weeks of age were purchased from the Jackson Laboratory (stock #000664 and stock #000651) were used for the preparation of BMDCs and for *in vivo* experiments. These mice were maintained in the animal facility at the Sloan Kettering Institute. All procedures were performed in strict accordance with the recommendations in the Guide for the Care and Use of Laboratory Animals of the National Institutes of Health. The protocol was approved by the Committee on the Ethics of Animal Experiments of Sloan Kettering Cancer Institute. STING^Gt/Gt^ mice were generated in the laboratory of Dr. Russell Vance (University of California, Berkeley). Foxp3^gfp^ and Foxp3^DTR^ mice were generated in the laboratory of Dr. Alexander Y. Rudensky (Memorial Sloan Kettering Cancer Center).

MMTV-PyMT mice were provided by Ming Li (Memorial Sloan Kettering Cancer Center). cGAS^-/-^ mice were generated in Herbert (Skip) Virgin’s laboratory (Washington University). STAT1^-/-^, STAT2^-/-^ and OX40^-/-^ were purchased from Jackson Laboratory. OX40^-/-^Foxp3^gfp^ mice were generated in our lab.

#### TIL isolation and flow cytometry

For TIL or myeloid cells analysis, tumors were minced prior to incubation with Liberase (1.67 Wünsch U/ml) and DNaseI (0.2 mg/ml) for 30 min at 37°C. Tumors were then homogenized by gentleMACS dissociator and filtered through a 70-μm nylon filter. Cell suspensions were washed and resuspended with complete RPMI. For cytokine production analysis, cells were restimulated with Cell Stimulation Cocktail (Thermo Fisher) and GolgiPlug (BD Biosciences) in complete RPMI for 6 h at 37°C. Cells were incubated with appropriate antibodies for surface labeling for 30 min at 4°C after staining dead cells with LIVE/DEAD™ Fixable Aqua Stain (Thermo Fisher). Cells were fixed and permeabilized using Foxp3 fixation and permeabilization kit (Thermo Fisher) for 1 hour at 4°C and then stained for Granzyme B, Foxp3, IFN*γ* and TNF*α*.

To analyze transgene expression, cells were infected with various viruses at a MOI of 10 or mock-infected. At 24 h post infection, cells were collected and the cell viability was determined by labeling with LIVE/DEAD™ Fixable Aqua Stain (Thermo Fisher) 15 min at 4°C. Cells were then sequentially stained with hFlt3L primary antibody, PE-conjugated goat-anti-mouse IgG antibody and AF647-conjugated anti-mOX40L antibody at 4°C, 15 min for each step.

For dendritic cell maturation assay, cells were infected with virus at a MOI of 10 and collected at 16 h post infection. Then cells were stained with anti-CD86 antibody for surface labeling for 30 min at 4 °C. LIVE/DEAD™ Fixable Aqua Stain (Thermo Fisher) was used to stain dead cells. Cells were analyzed using the BD LSRFortessa flow cytometer (BD Biosciences). Data were analyzed with FlowJo software (Treestar).

#### RNA isolation and Real-time PCR

For the generation of BMDCs, the bone marrow cells (5 million cells in each 15 cm cell culture dish) were cultured in RPMI-1640 medium supplemented with 10% fetal bovine serum (FBS) in the presence of 30 ng/ml GM-CSF (BioLegend) for 10-12 days.

To generate human monocyte-derived dendritic cells, peripheral blood mononuclear cells (PBMC) were prepared by centrifugation on a Ficoll gradient. Monocytes layer was collected and plated to tissue culture dish. After 1 h, non-adherent cells were washed off. The remaining cells were cultured for 5-7 days in RPMI-1640 supplemented with antibiotics (penicillin and streptomycin) and 10% FCS in the presence of 1000 IU/ml GM-CSF (PeproTech) and 500 IU/ml IL-4 (PeproTech).

Cells were infected with various viruses at a MOI of 10 for 1 hour or mock-infected. The inoculum was removed, and the cells were washed with PBS twice and incubated with fresh medium. RNA was extracted from whole-cell lysates with RNeasy Plus Mini kit (Qiagen) and was reverse-transcribed with cDNA synthesis kit (Thermo Fisher). Real-time PCR was performed in triplicate with SYBR Green PCR Master Mix (Life Technologies) and Applied Biosystems 7500 Real-time PCR Instrument (Life Technologies) using gene-specific primers. Relative expression was normalized to the levels of glyceraldehyde-3-phosphate dehydrogenase (GAPDH). The primer sequences for quantitative real-time PCR are listed in Table S1.

#### Tumor challenge and treatment

For tumor immune cells analysis, B16-F10 cells were implanted intradermally into right and left flanks of the mice (5*×*10^5^ to the right flank and 2.5*×*10^5^ to the left flank). At 7 to 9 days after implantation, the tumors at the right flank were injected with 4*×*10^7^ PFU of rMVA, MVA*Δ*E5R or PBS twice, 2 or 3 days apart. Tumors, spleens and/or tumor draining lymph nodes were harvested two days after second injection. In some experiments, 50 μg of *α*IFNAR-1 antibody (MAR1-5A3, BioXcell) were injected into the tumors together with rMVA.

For survival experiments, 2*×*10^5^ B16-F10 cells were implanted intradermally into the shaved skin on the right flank of WT C57BL/6J mice or age-matched cGAS^-/-^, STING^Gt/Gt^, STAT2^-/-^, STAT1^-/-^ mice. In some experiments, 2*×*10^5^ A20 cells were implanted intradermally into the right flank of WT BALB/cJ mice. At 6 to 9 days after implantation, tumor sizes were measured and tumors that are 3 mm in diameter or larger were injected with 4*×*10^7^ PFU of rMVA or PBS when the mice were under anesthesia. Viruses were injected twice weekly as specified in each experiment and tumor sizes were measured twice a week. Tumor volumes were calculated according to the following formula: l (length) × w (width) × h (height)/2. Mice were euthanized for signs of distress or when the diameter of the tumor reached 10 mm. For depletion of T cells, depletion antibodies for CD8^+^ and CD4^+^ cells (200 μg of clone 2.43 and GK1.5, BioXcell) were injected intraperitoneally twice weekly starting 1 day before viral injection, and they were used until the animals either died, were euthanized, or were completely clear of tumors. In some experiments, 25 μg of FTY720 diluted in 100 μl deionized water was injected intraperitoneally twice weekly starting 1 day before viral injection.

For depletion of Tregs, 2*×*10^5^ B16-F10 cells were implanted intradermally into the shaved skin on the right flank of Foxp3^DTR^ mice. Diphtheria toxin (DT) were injected were injected intraperitoneally twice weekly starting 1 day before viral injection, and they were used until the endpoint. In the bilateral tumor implantation model, B16-F10 cells were implanted intradermally into right and left flanks of C57BL/6J mice (5*×*10^5^ to the right flank and 1*×*10^5^ to the left flank). At 7 days after implantation, the tumors at the right flank were injected with 4*×*10^7^ PFU of rMVA or PBS. 250μg *α*PD-L1 antibody (10F.9G2, BioXcell) was injected intraperitoneally twice weekly.

For the tumor rechallenge study, the survived mice (more than 40 days after initiation of intratumoral virotherapy) were rechallenged with intradermal delivery of a lethal dose of B16-F10 (1*×*10^5^ cells) at the contralateral side.

#### ELISpot assay

Spleens were mechanically disrupted by gentleMACS^TM^ dissociator and red blood cells were lysed by ACK lysing buffer. 1*×*10^6^ splenocytes were co-cultured with 2.5*×*10^5^ irradiated B16-F10 in complete RPMI medium overnight. IFN*γ*^+^ splenocytes were detected by Mouse IFN*γ* ELISPOT kit (BD Biosciences)

#### In vitro Treg suppression assay

5*×*10^5^ WT B16-F10 cells were implanted intradermally into the right and left flanks of Foxp3^gfp^ and OX40^-/-^Foxp3^gfp^ mice. Tumors and spleens were harvested when tumor sizes reached 5 mm in diameter or larger and processed into single cell suspensions as described above. Cells were stained with anti-CD45.2 (AF700), CD4 (Pacific Blue) and OX40 (PE) antibodies for 30 min at 4 °C. LIVE/DEAD™ Fixable Aqua Stain (Thermo Fisher) was used to stain dead cells. Treg Cells were sorted into CD4^+^GFP^+^OX40^hi^ and CD4^+^GFP^+^OX40^lo^ populations. Naive CD4^+^ T Cells were sorted from mouse spleen. Spleens from CD45.1 congenic mouse were harvested, chopped and digested with Collagenase D (2.5 mg/ml) and DNaseI (50 μg/ml) for 30 min at 37°C. CD11c^+^ DCs were then isolated by CD11c MicroBeads (Miltenyi Biotec). CTV-labeled 4*×*10^4^ native CD4^+^ T cells were co-cultured with 1*×*10^5^ CD11c^+^ DCs. Purified CD4^+^GFP^+^OX40^hi^, CD4^+^GFP^+^OX40^lo^ or CD4^+^GFP^+^OX40^-/-^ cells were seeded in indicated ratios and cultured in complete RPMI supplemented with 1 μg/ml anti-CD3 antibody for 3 days. Cell were then collected and stained with APC anti-CD4 antibody for 30 min at 4 °C after staining dead cells with Zombie Nir. Data were acquired using the BD LSRFortessa flow cytometer (BD Biosciences). Data were analyzed with FlowJo software (Tree Star).

#### Bulk tumor RNA-seq

5*×*10^5^ WT B16-F10 cells were implanted intradermally into right and left flanks of WT C57BL/6J or age-matched STING^Gt/Gt^ mice. At 7 days after implantation, the tumors were injected with 4*×*10^7^ PFU of rMVA or PBS. Tumors were harvested 1 day after injection and processed into single cell suspension as described previously. RNA was extracted from whole-cell lysates with RNeasy Plus Mini kit (Qiagen, Hilden, Germany) for RNA-seq.

Library prep and sequencing: Following RNA isolation, total RNA integrity was checked using a 2100 Bioanalyzer (Agilent Technologies, Santa Clara, CA). RNA concentrations were measured using the NanoDrop system (Thermo Fisher Scientific, Inc., Waltham, MA). Preparation of RNA sample library and RNA-seq were performed by the Genomics Core Laboratory at Weill Cornell Medicine. Messenger RNA was prepared using TruSeq Stranded mRNA Sample Library Preparation kit (Illumina, San Diego, CA), according to the manufacturer’s instructions. The normalized cDNA libraries were pooled and sequenced on Illumina NovaSeq6000 sequencer with pairend 50 cycles.

#### Treg RNA-seq

5*×*10^5^ WT B16-F10 cells were implanted intradermally into right and left flanks of Foxp3^GFP^ mice. When tumor diameter reached 5 mm, mice were euthanized, and tumors and spleens were harvested and processed into single cell suspension as described before. CD4^+^GFP^+^OX40^hi^ and CD4^+^GFP^+^OX40^low^ populations from tumors and spleens were purified by FACS sorting. RNA was extracted with RNeasy Plus Micro kit (Qiagen). Following RNA isolation, total RNA integrity was checked using a 2100 Bioanalyzer (Agilent Technologies, Santa Clara, CA). RNA concentrations were measured using the NanoDrop system (Thermo Fisher Scientific, Inc., Waltham, MA). The cDNA synthesis and amplification were performed by SMART-Seq v4 ultra low input RNA kit (Takara Bio USA, Mountain View, CA, USA) starting with less than 1 ng of total RNA from each sample. 150 pg of qualified full-length double strand cDNA was used and processed to Illumina library construction with the Nextera XT DNA Library Preparation Kits (Illumina, San Diego, CA). Then the normalized cDNA libraries were pooled and sequenced on Illumina NovaSeq6000 sequencer with pair-end 50 cycles.

#### RNA-seq data analysis

The raw sequencing reads in BCL format were processed through bcl2fastq 2.19 (Illumina) for FASTQ conversion and demultiplexing. After trimming the adaptors with cutadapt (version1.18)( https://cutadapt.readthedocs.io/en/v1.18/), RNA reads were aligned and mapped to the GRCh38 human reference genome by STAR (Version2.5.2) (https://github.com/alexdobin/STAR) (Dobin et al., 2013), and transcriptome reconstruction was performed by Cufflinks (Version 2.1.1) (http://cole-trapnell-lab.github.io/cufflinks/). The abundance of transcripts was measured with Cufflinks in Fragments Per Kilobase of exon model per Million mapped reads (FPKM) (Trapnell et al., 2013; Trapnell et al., 2010). Gene expression profiles were constructed for differential expression, cluster, and principle component analyses with the DESeq2 package (https://bioconductor.org/packages/re-lease/bioc/html/DESeq2.html) (Love et al., 2014). For differential expression analysis, pairwise comparisons between two or more groups using parametric tests where read-counts follow a negative binomial distribution with a gene-specific dispersion parameter. Corrected p-values were calculated based on the Benjamini-Hochberg method to adjusted for multiple testing.

The GSEA analysis was done using GSEA software version 4.0.3 (Subramanian et al., 2005) from the Broad Institute(http://www.gsea-msigdb.org/gsea/index.jsp), which uses predefined gene sets from the Molecular Signatures Database (MSigDB v7.4) (Subramanian et al., 2005). We used the hallmark gene sets collection for the present study. Genes were ranked by the test statistic value obtained from differential expression analysis and the pre-ranked version of the tool was used to identify significantly enriched biological pathways. The minimum and maximum criteria for selection of gene sets from the collection were 15 and 500 genes, respectively.

#### Human tumor specimens

Fresh biopsy samples from patients with Extramammary Paget’s disease were obtained at the dermatology service in the Department of Medicine of Memorial Sloan Kettering Cancer Center. Written informed consents were obtained from patients enrolled in the protocol approved by Memorial Sloan Kettering Cancer Center Institutional Review Board (IRB). Studies were conducted in accordance with National Institutes of Health and institutional guidelines for human subject research. Tumor tissues were cut into small pieces using a pair of fine scissors. They were infected with rhMVA or mock-infected. Cells were collected after 24 h and processed for FACS analyses.

#### Statistical analysis

Two-tailed unpaired Student’s t test was used for comparisons of two groups in the studies. Survival data were analyzed by log-rank (Mantel-Cox) test. The p values deemed significant are indicated in the figures as follows: *, p < 0.05; **, p < 0.01; ***, p < 0.001; ****, p < 0.0001. The numbers of animals included in the study are discussed in each figure legend.

## Acknowledgements

We thank Alexander Y. Rudensky for Foxp3^gfp^ and Foxp3^DTR^ mice. We thank the Flow Cytometry Core Facility and Molecular Cytology Core Facility at the Sloan Kettering Institute and Genomic Resources Core Facility at Weill Cornell Medical College. This work was supported by NIH grants K-08 AI073736 (L.D.), R56AI095692 (L.D.), R03 AR068118 (L.D.), R01 CA56821 (J.D.W). R01 AI091707 (C.M.R.), Charles H. Revson Senior Fellowship in Biomedical Science (J.M.L), a Black Family Metastasis Center Fellowship (J.M.L.), Society of Memorial Sloan Kettering (MSK) research grant (L.D.), MSK Technology Development Fund (L.D.), Parker Institute for Cancer Immunotherapy Career Development Award (L.D.), sponsored research agreement from IMVAQ Therapeutics (L.D.). This work was supported in part by the Swim across America (J.D.W., T.M.), Ludwig Institute for Cancer Research (J.D.W., T.M.), This research was also funded in part through the NIH/NCI Cancer Center Support Grant P30 CA008748.

## Author Contributions

Author contributions: N.Y., Y.W. and L.D. were involved in all aspect of this study, including conceiving the project, designing and performing experiments, data analyses and interpretation, and manuscript writing. S.L. assisted some mouse experiments and human tumor ex vivo infection experiments and analyzed the data. G.M, J.W., W.Y. J.C. assisted with construct designs and viral engineering. J.M.L. analyzed RNA-seq data on regulatory T cells. A.Y.T. and T.Z. analyzed the bulk RNA-seq data from tumors. A.R. provided human tumor samples. J.D.W., T.M, C.M.R. and J.Z.X. assisted in experimental design, data interpretation, and manuscript preparation. All authors are involved in manuscript preparation. L.D. provided overall supervision of the study.

