## Supplementary figures and table for "Intratumoral delivery of engineered recombinant modified vaccinia virus Ankara expressing Flt3L and OX40L generates potent antitumor immunity through activating the cGAS/STING pathway and depleting tumor-infiltrating regulatory T cells"

This file contains:

- Supplementary Figure 1
- Supplementary Figure 2
- Supplementary Figure 3
- Supplementary Figure 4
- Supplementary Figure 5
- Supplementary Figure 6
- Supplementary Table 1

### Figure S1

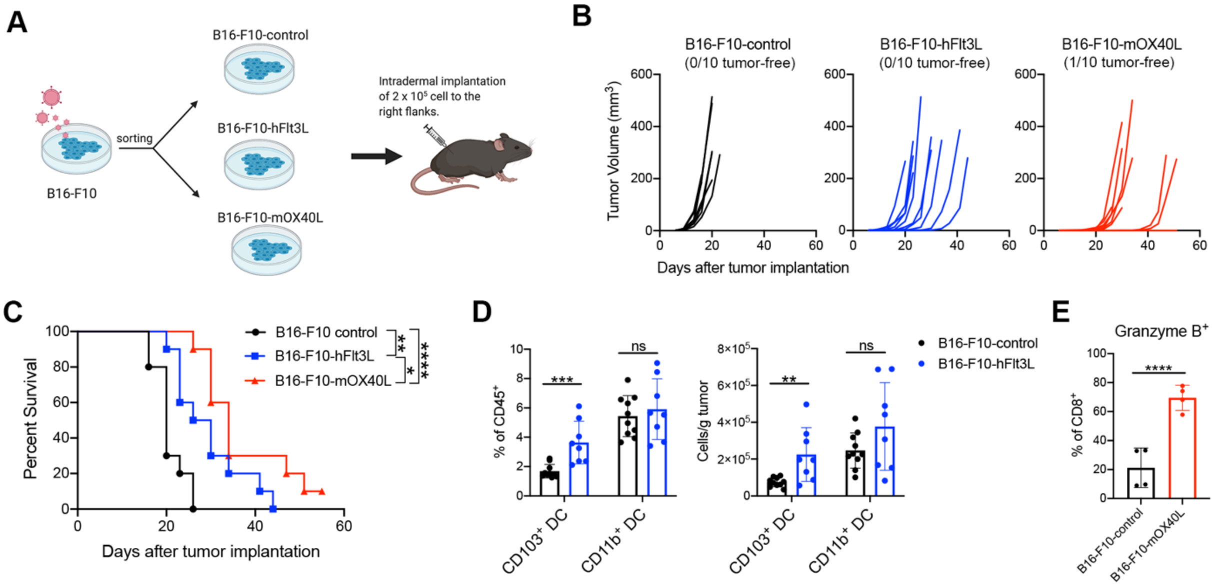

A
B16-F10-control
B16-F10-hFlt3L
B16-F10-mOX40L
B16-F10
sorting
Intradermal implantation of 2 x 10⁵ cell to the right flanks.
B
B16-F10-control (0/10 tumor-free)
B16-F10-hFlt3L (0/10 tumor-free)
B16-F10-mOX40L (1/10 tumor-free)
Tumor Volume (mm³)
Days after tumor implantation
C
B16-F10 control
B16-F10-hFlt3L
B16-F10-mOX40L
Percent Survival
Days after tumor implantation
D
% of CD45⁺
Cells/g tumor
CD103⁺ DC
CD11b⁺ DC
B16-F10-control
B16-F10-hFlt3L
E
Granzyme B⁺
% of CD8⁺
B16-F10-control
B16-F10-mOX40L

**Figure S1.** Overexpression of human Flt3L or murine OX40L on B16-F10 tumor cells enhances immunogenicity of the tumors.
(A) B16-F10 were transduced with retrovirus to generate hFlt3L or mOX40L-expressing stable cell lines. C57BL/6J mice were intradermally implanted with 2 x $10^5$ B16-F10-hFlt3L, B16-F10-mOX40L or B16-F10 control cells.
(B) Tumor growth curve (*n=10*).
(C) Kaplan-Meier survival curve (*n=10; *P < 0.05, **P < 0.01, ****P < 0.0001, Mantel-Cox test*).
(D) Percentages and absolutes number of $CD103^+$ DCs and $CD11b^+$ DCs in B16-F10-hFlt3L or B16-F10-control tumors. Data are means ± SD (*n=8 or 10; **P < 0.01, ***P < 0.001, t test)*.
(E) Percentages Granzyme $B^+$ $CD8^+$ and Granzyme $B^+$ $CD4^+$ in B16-F10-hFlt3L or B16-F10-control tumors. Data are means ± SD (*n=4; ****P < 0.0001, t test)*.

### Figure S2

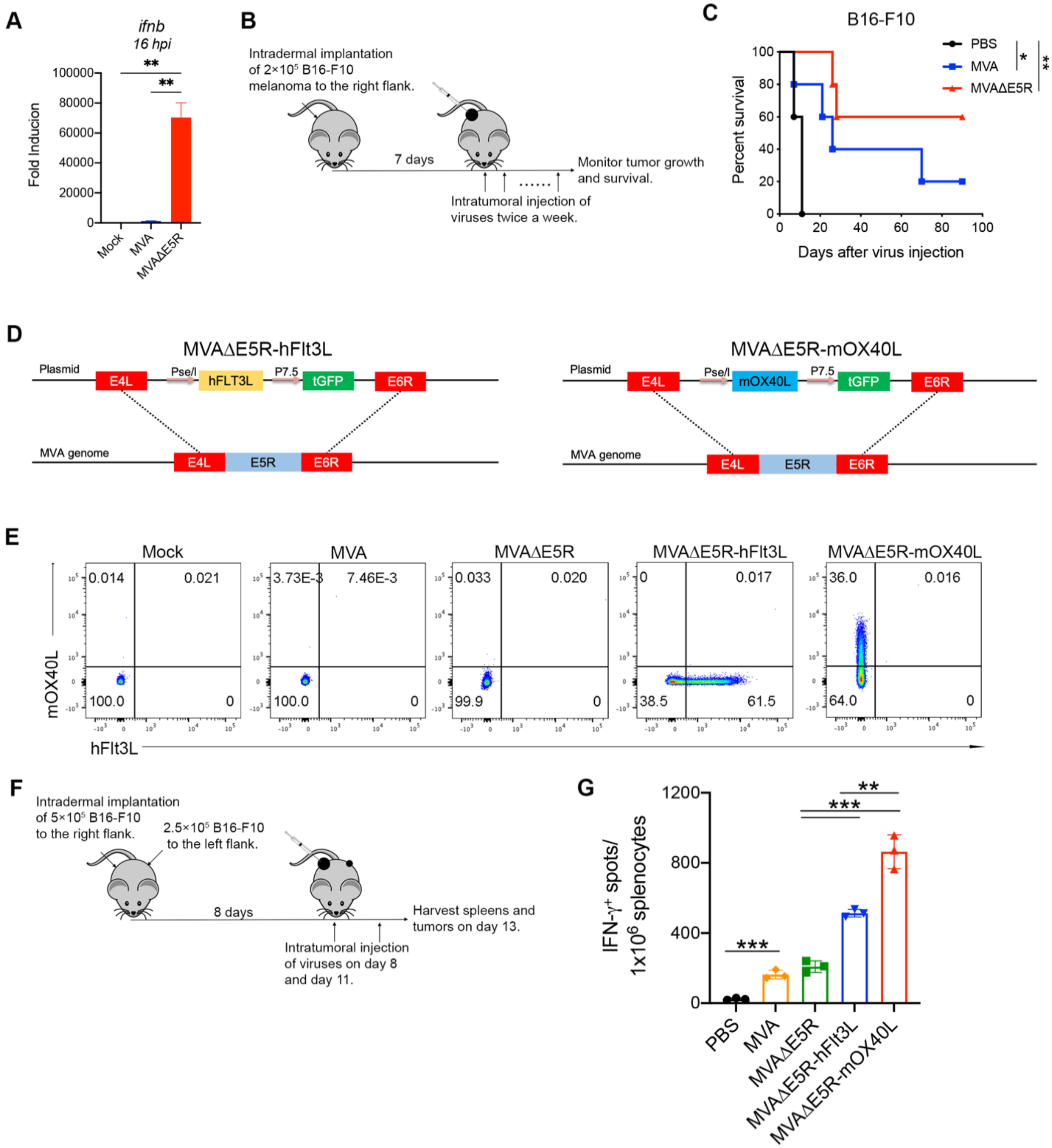

A
ifnb
16 hpi
Fold Inducion
Mock
MVA
MVAΔE5R
B
Intradermal implantation of 2×10⁵ B16-F10 melanoma to the right flank.
7 days
Monitor tumor growth and survival.
Intratumoral injection of viruses twice a week.
C
B16-F10
PBS
MVA
MVAΔE5R
Percent survival
Days after virus injection
D
MVAΔE5R-hFlt3L
Plasmid
E4L
Pse/I
hFLT3L
P7.5
tGFP
E6R
MVA genome
E5R
MVAΔE5R-mOX40L
mOX40L
E
Mock
MVA
MVAΔE5R
MVAΔE5R-hFlt3L
MVAΔE5R-mOX40L
mOX40L
hFlt3L
F
Intradermal implantation of 5×10⁵ B16-F10 to the right flank.
2.5×10⁵ B16-F10 to the left flank.
8 days
Harvest spleens and tumors on day 13.
Intratumoral injection of viruses on day 8 and day 11.
G
IFN-γ⁺ spots/ 1x10⁶ splenocytes
PBS
MVA
MVAΔE5R
MVAΔE5R-hFlt3L
MVAΔE5R-mOX40L

**Figure S2.** Incremental engineering of MVA with deletion of E5R gene and expression of human Flt3L or murine OX40L improves antitumor effects.
(A) Relative mRNA expression levels of *Ifnb* in BMDCs infected with MVA or MVAΔE5R. Data are means ± SD (*n=3; **P < 0.01, t test).*
(B) Schematic diagram of IT MVA, MVAΔE5R or PBS in a unilateral B16-F10 melanoma implantation model.
(C)Kaplan-Meier survival curve of mice treated with MVA, MVAΔE5R or PBS in a unilateral B16-F10 implantation model (*n=10; *P < 0.05, **P < 0.01, Mantel-Cox test*).
(D) Schematic diagrams for the generation of MVAΔE5R-hFlt3L or MVAΔE5R-mOX40L through homologous recombination.
(D) Representative flow cytometry plots of expression of hFlt3L or mOX40L by MVA, MVAΔE5R, MVAΔE5R-hFlt3L, MVAΔE5R-mOX40L or mock-infected BHK21 cells.
(E) Representative flow cytometry plots of expression of hFlt3L or mOX40L by MVA, MVAΔE5R, MVAΔE5R-hFlt3L, MVAΔE5R-mOX40L or mock-infected BHK21 cells.
(F) Schematic diagram of IT MVA, MVAΔE5R, MVAΔE5R-hFlt3L, MVAΔE5R-mOX40L or PBS in a bilateral B16-F10 melanoma implantation model.
(G) IFN-$\gamma^+$ splenocytes from MVA, MVAΔE5R, MVAΔE5R-hFlt3L, MVAΔE5R-mOX40L or PBS-treated mice. Data are means ± SD (*n=3; *P < 0.05, ***P < 0.001, t test).*

### Figure S3

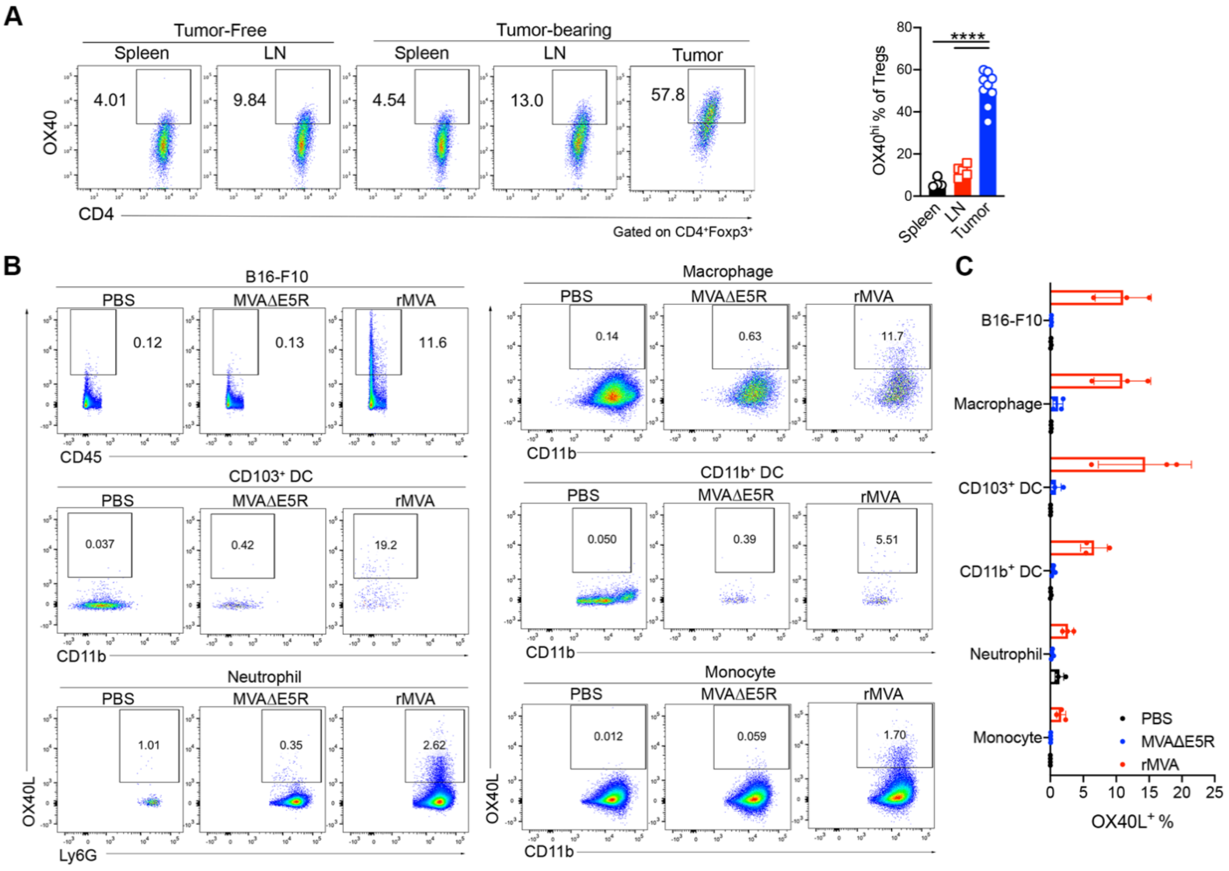
A
Tumor-Free
Tumor-bearing
Spleen
LN
Tumor
4.01
9.84
4.54
13.0
57.8
OX40
CD4
Gated on CD4⁺Foxp3⁺
OX40hi % of Tregs
****
B
B16-F10
PBS
MVAΔE5R
rMVA
0.12
0.13
11.6
CD45
CD103⁺ DC
0.037
0.42
19.2
CD11b
Neutrophil
1.01
0.35
2.62
Ly6G
OX40L
Macrophage
0.14
0.63
11.7
CD11b⁺ DC
0.050
0.39
5.51
Monocyte
0.012
0.059
1.70
C
B16-F10
Macrophage
CD103⁺ DC
CD11b⁺ DC
Neutrophil
Monocyte
PBS
MVAΔE5R
rMVA
OX40L⁺ %

**Figure S3.** OX40 expression on T cells in lymphoid organs and in tumors and OX40L expression in tumors and tumor-infiltrating cells after IT rMVA.
(A) Representative flow cytometry plots of OX40 expression on $CD4^{+}Foxp3^{+}$ T cells in the spleens, lymph nodes or tumors from naïve or B16-F10 tumor-bearing mice.
(B) Representative flow cytometry plots of OX40L expression on B16-F10 tumor cells or myeloid cells in the tumors injected with MVAΔE5R, rMVA or PBS as control.
(C) Percentages of $OX40L^{+}$ B16-F10 cells or myeloid cells in the tumors. Data are means ± SD (*n=3~5)*.

**Figure S4**

**A**

**OX40hi Tumor versus spleen**
Chemokines & receptors

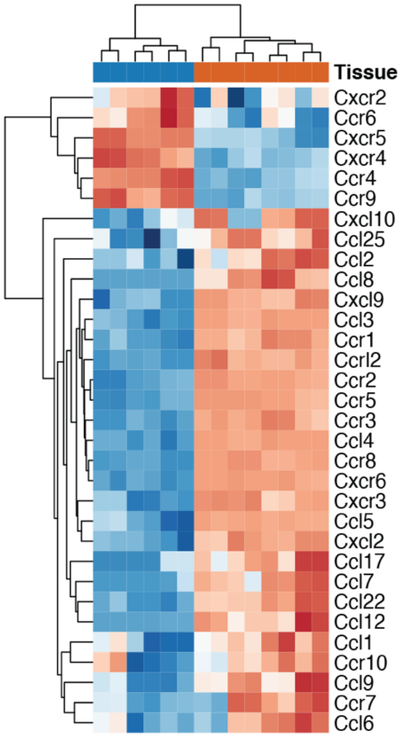


**B**

**OX40hi Tumor versus spleen**
Immune suppressors

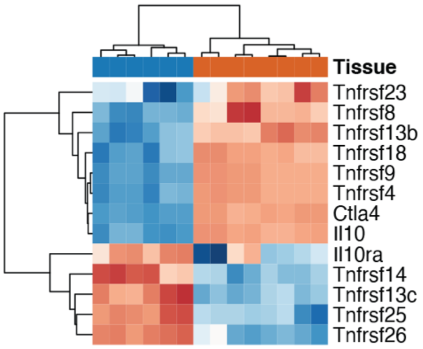


**C**

**OX40hi Tumor versus spleen**
GO Oxidative Phosphorylation

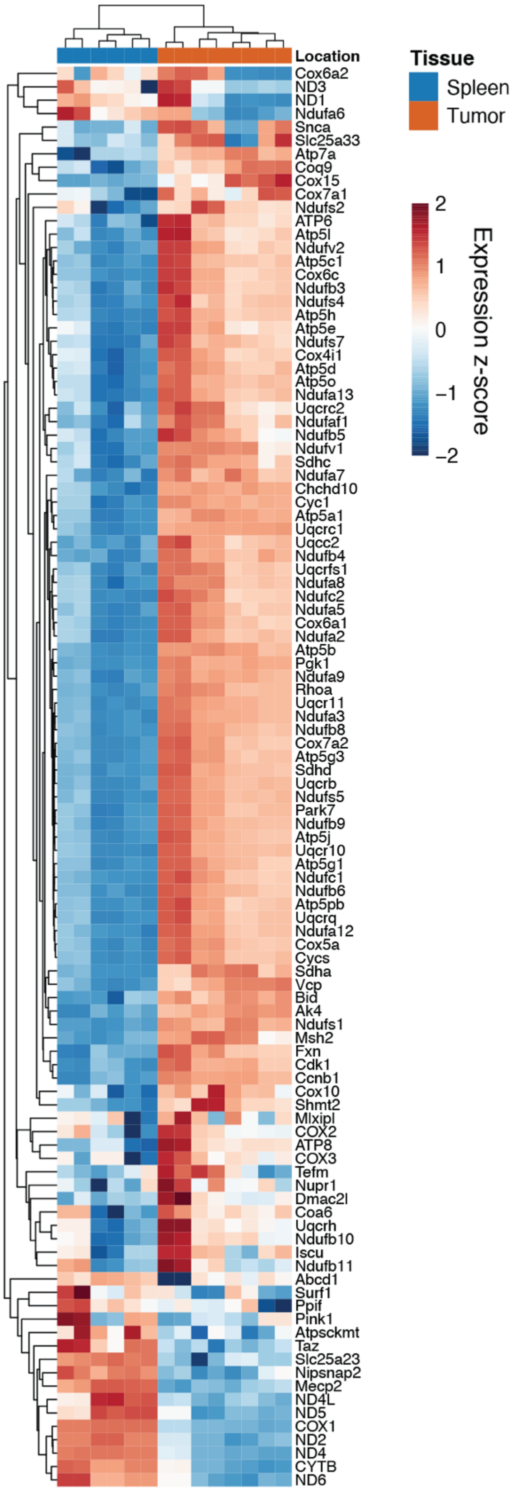

**Figure S4.** Heatmaps of differential gene expression in OX40$^{hi}$ Tregs isolated from tumors vs. those from spleens.
(A) Differential gene expression of chemokines and chemokine receptors
(B) Differential gene expression of immune suppressive genes
(C) Differential gene expression of genes involved in oxidative phosphorylation.

**Figure S5**

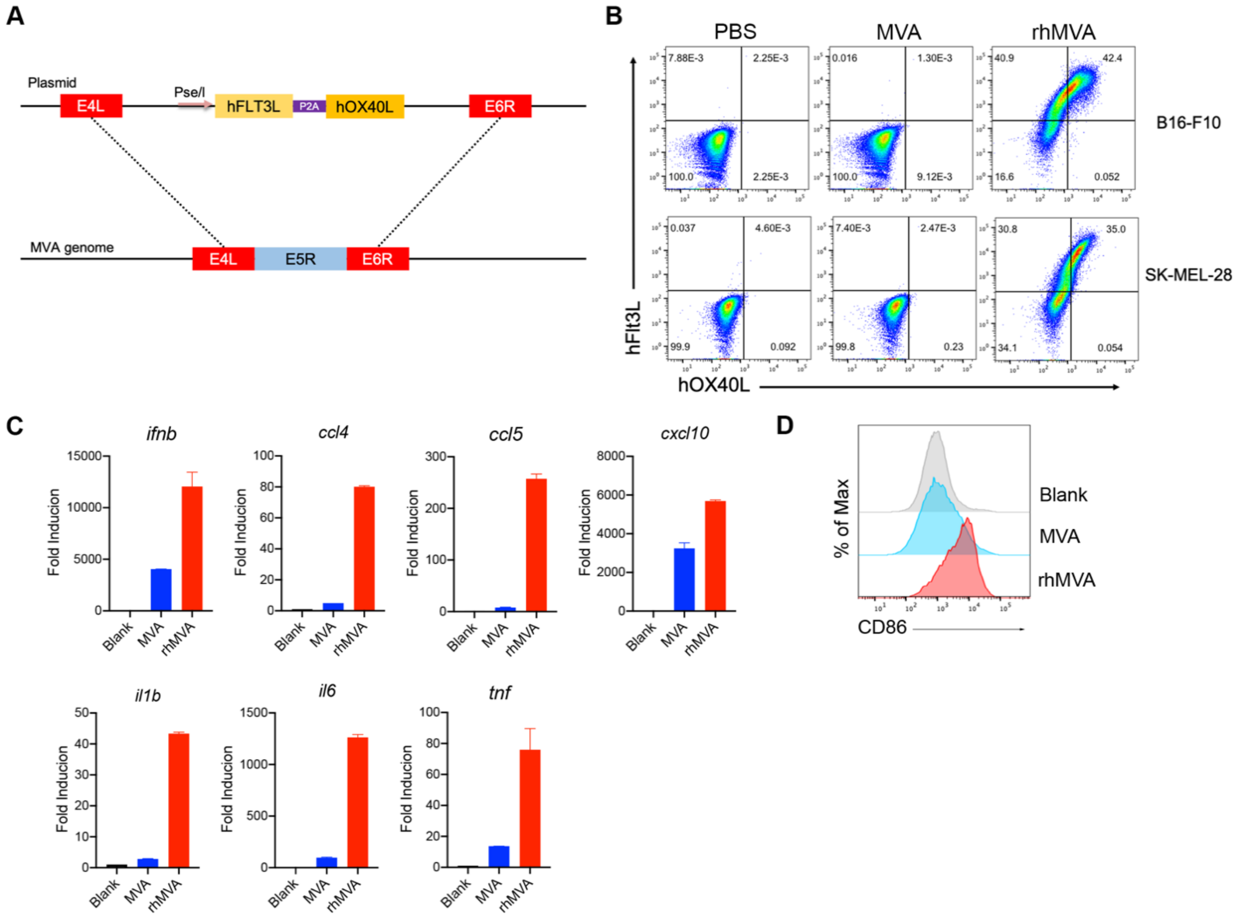
A
Plasmid
E4L
Pse/l
hFLT3L
P2A
hOX40L
E6R
MVA genome
E4L
E5R
E6R
B
PBS
MVA
rhMVA
B16-F10
SK-MEL-28
hFlt3L
hOX40L
C
ifnb
ccl4
ccl5
cxcl10
il1b
il6
tnf
Fold Inducion
Blank
MVA
rhMVA
D
% of Max
Blank
MVA
rhMVA
CD86

**Figure S5. Clinical candidate rhMVA induces innate immunity and promotes maturation of human monocyte-derived DCs (moDCs).**

(A) Schematic diagram for the generation of rhMVA through homologous recombination.

(B) Representative flow cytometry plots of expression of hFlt3L or hOX40L by rMVA-infected B16-F10 cells and SK-MEL-28 cells.

(C) Relative mRNA expression levels of *ifnb*, *ccl4*, *ccl5*, *cxcl10*, *il1b*, *il6* and *tnf* in moDCs infected with MVA or rhMVA.

(D) Mean fluorescence intensity of CD86 expressed by MoDCs infected with MVA or rhMVA.

#### Figure S6

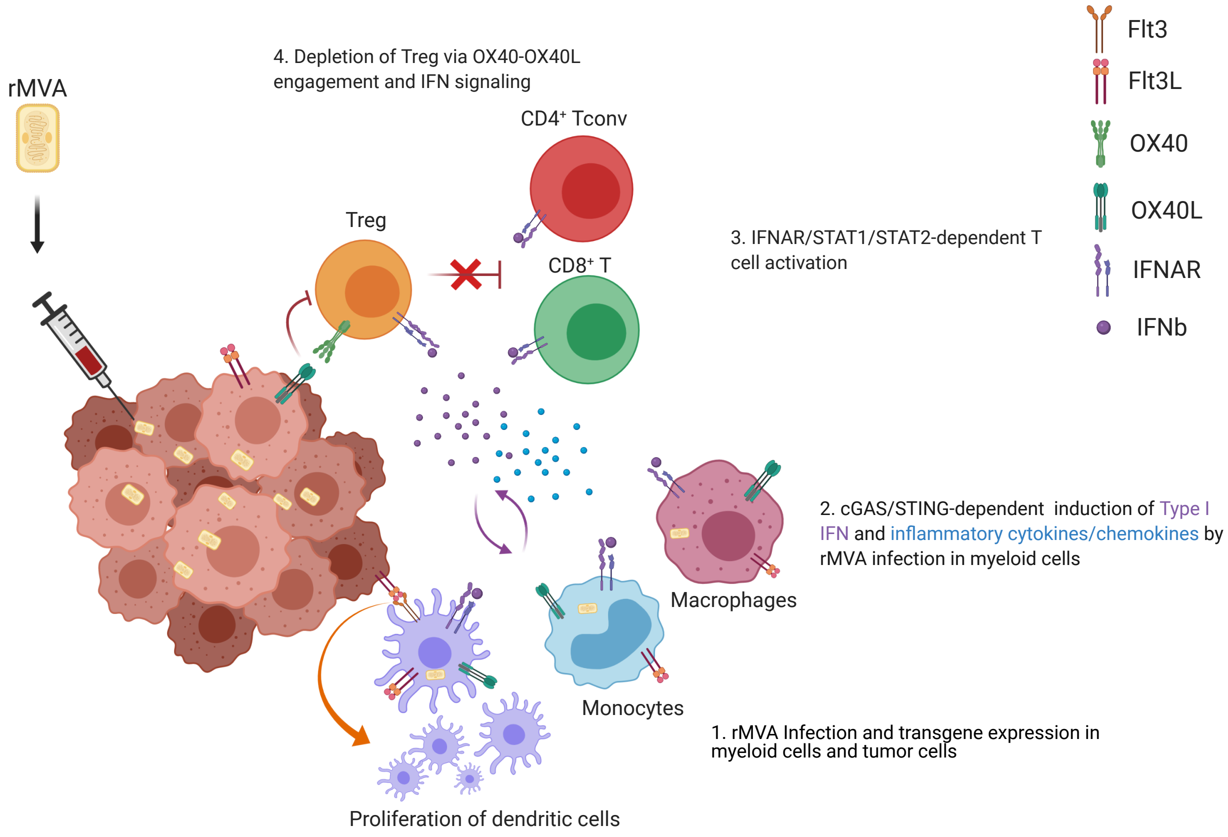
rMVA
4. Depletion of Treg via OX40-OX40L engagement and IFN signaling
CD4⁺ Tconv
Treg
CD8⁺ T
3. IFNAR/STAT1/STAT2-dependent T cell activation
Flt3
Flt3L
OX40
OX40L
IFNAR
IFNb
2. cGAS/STING-dependent induction of Type I IFN and inflammatory cytokines/chemokines by rMVA infection in myeloid cells
Macrophages
Monocytes
1. rMVA Infection and transgene expression in myeloid cells and tumor cells
Proliferation of dendritic cells

**Figure S6. Working model.** IT injection of rMVA results in the infection of tumor-infiltrating myeloid cells, including macrophages, monocytes, and dendritic cells, as well as tumor cells. This leads to the activation of cGAS/STING-mediated cytosolic DNA-sensing pathway and the production of type I IFN and cytokines and chemokines that are important for $CD8^+$ and $CD4^+$ T cell proliferation and activation (as indicated by Granzyme B, TNF, and IFN-γ expression). Flt3L expression of the tumor microenvironment facilitates the proliferation of $CD103^+$ DCs in the tumors. OX40L expression by myeloid cell populations and tumor cells results in the depletion of $OX40^{hi}$ Tregs infiltrating the tumors via OX40L-OX40 ligation, which is promoted by type I IFN. This leads to the blunting of their inhibition on tumor-specific effector $CD4^+$ and $CD8^+$ T cells. Taken together, IT delivery of rMVA results in the alteration of tumor immunosuppressive microenvironment through activation of innate immunity and boosting of antitumor T cells by depletion of $OX40^{hi}$ regulatory T cells.

**Table S1**. Primers for Real-time PCR

| Species | Gene | Direction | Sequence |
|---|---|---|---|
| Mouse | ccl4 | Forward | 5'-GCCCTCTCTCTCCTCTTGCT-3' |
| | ccl4 | Reverse | 5'-CTGGTCTCATAGTAATCCATC-3' |
| | ccl5 | Forward | 5'-GCCCACGTCAAGGAGTATTTCTA-3' |
| | ccl5 | Reverse | 5'-ACACACTTGGCGGTTCCTTC-3' |
| | cxcl10 | Forward | 5'-GTCAGGTTGCCTCTGTCTCA-3' |
| | cxcl10 | Reverse | 5'-TCAGGGAAGAGTCTGGAAAG-3' |
| | cxcl9 | Forward | 5'-GGAACCCTAGTGATAAGGAATGCA-3' |
| | cxcl9 | Reverse | 5'-TGAGGTCTTTGAGGGATTTGTAGTG-3' |
| | ifna | Forward | 5'-CCTGTGTGATGCAGGAACC-3' |
| | ifna | Reverse | 5'-TCACCTCCCAGGCACAGA-3' |
| | ifnb | Forward | 5'-TGGAGATGACGGAGAAGATG-3' |
| | ifnb | Reverse | 5'-TTGGATGGCAAAGGCAGT-3' |
| | GAPDH | Forward | 5'-ATCAAGAAGGTGGTGAAGCA-3' |
| | GAPDH | Reverse | 5'-AGACAACCTGGTCCTCAGTGT-3' |
| | ccl4 | Forward | 5'- AAAACCTCTTTGCCACCAATACC-3' |
| | ccl4 | Reverse | 5'- GAGAGCAGAAGGCAGCTACTAG-3' |
| Human | cxcl10 | Forward | 5'-ATTTGCTGCCTTATCTTTCTG-3' |
| | cxcl10 | Reverse | 5'-TCTCACCCTTCTTTTTCATTGTAG-3' |
| | ifnb | Forward | 5'-GCACTGGCTGGAATGAGACT-3' |
| | ifnb | Reverse | 5'-CCTTGGCCTTCAGGTAATG-3' |
| | il6 | Forward | 5'-AATTCGGTACATCCTCGACGG-3' |
| | il6 | Reverse | 5'-TTGGAAGGTTCAGGTTGTTTTCT-3' |
| | tnf | Forward | 5'-AATAGGCTGTTCCCATGTAGC-3' |
| | tnf | Reverse | 5'-AGAGGCTCAGCAATGAGTGA-3' |
| | GAPDH | Forward | 5'-ATCAAGAAGGTGGTGAAGCA-3' |
| | GAPDH | Reverse | 5'-GTCGCTGTTGAAGTCAGAGGA-3' |
